# ORANGE: A CRISPR/Cas9-based genome editing toolbox for epitope tagging of endogenous proteins in neurons

**DOI:** 10.1101/700187

**Authors:** Jelmer Willems, Arthur P.H. de Jong, Nicky Scheefhals, Harold D. MacGillavry

**Author notes:** CONTACT INFO Correspondence to: H.D.M.

## Abstract

The correct subcellular distribution of protein complexes establishes the complex morphology of neurons and is fundamental to their functioning. Thus, determining the dynamic distribution of proteins is essential to understand neuronal processes. Fluorescence imaging, in particular super-resolution microscopy, has become invaluable to investigate subcellular protein distribution. However, these approaches suffer from the limited ability to efficiently and reliably label endogenous proteins. We developed ORANGE: an Open Resource for the Application of Neuronal Genome Editing, that mediates targeted genomic integration of fluorescent tags in neurons. This toolbox includes a knock-in library for in-depth investigation of endogenous protein distribution, and a detailed protocol explaining how knock-in can be developed for novel targets. In combination with super-resolution microscopy, ORANGE revealed the dynamic nanoscale organization of endogenous neuronal signaling molecules, synaptic scaffolding proteins, and neurotransmitter receptors. Thus, ORANGE enables quantitation of expression and distribution for virtually any protein in neurons at high resolution and will significantly further our understanding of neuronal cell biology.

## INTRODUCTION

Neurons are highly complex cells with numerous functionally and structurally distinct subcellular compartments that are each composed of unique repertoires of molecular components. The correct targeting and localization of protein complexes, and their spatial organization within subcellular domains underlies virtually every aspect of neuronal functioning. Thus, investigating the dynamic distribution of proteins in neurons is critical for a mechanistic understanding of brain function. Precise localization of individual protein species using fluorescence microscopy has become an essential technique in many fields of neuroscience, and in particular for studies on synaptic function, where protein mislocalization at scales less than one micrometer can already have detrimental effects on synaptic efficacy (1). Recently developed super-resolution imaging methods now routinely achieve spatial resolution to only tens of nanometers, allowing determination of protein distributions at the molecular scale (2, 3). Consequently, these methods are highly sensitive to experimental alterations that affect protein organization, and efficient labeling techniques that accurately report the localization of endogenous proteins are critical.

Visualization of subcellular protein localization typically relies on antibody-based labeling approaches or overexpression of fluorescently tagged proteins, but both techniques have serious limitations (4). Immunocytochemistry largely relies on the availability of specific antibodies, which has severely hampered progress for many targets. For intracellular proteins, immuno-labeling requires fixation and permeabilization, manipulations that often introduce ultrastructural changes, interfere with protein localization, and preclude labeling and visualization of protein dynamics in live cells. Also, it is often desirable to sparsely label individual cells to measure protein distribution at high contrast, which is difficult to achieve with immunocytochemistry-based techniques. Expression of fluorophore-tagged proteins overcomes many of these issues, but exogenous introduction of recombinant proteins can lead to mislocalization or induce severe morphological and/or physiological effects. For instance, the overexpression of synaptic proteins such as PSD95 and Shank have pronounced effects on synapse number, content, structure, and physiology (5–8). Exorbitant expression levels can be circumvented by a replacement strategy, whereby a tagged protein is expressed on a knockdown or knockout background (9), but this will at best only approach endogenous levels, and is uncoupled from endogenous transcriptional or translational regulatory mechanisms. Recombinant antibody-based approaches have been developed for live-cell imaging of synaptic proteins, but have so far been restricted to a few targets (10–13). Furthermore, the generation and maintenance of transgenic animals is costly and time consuming, making it an inefficient approach for high-throughput tagging of neuronal proteins. Also, generally, transgenic labeling leads to global expression of tagged proteins in all cells, preventing imaging in distinct, isolated cells.

In view of the limitations of current techniques, we sought to develop a protein visualization strategy that meets the following criteria: 1) accurately reports a single protein species at endogenous levels and spatiotemporal expression; 2) does not interfere with protein localization and function; 3) is compatible with (super-resolution) light microscopy of live as well as fixed tissues; 4) is easily applied in both primary neuronal cultures and organotypic slices; 5) can be rapidly developed and expanded to many independent proteins of interest; and 6) allows for sparse labeling of neurons. We reasoned that labeling of endogenous proteins with fluorescent tags using targeted gene-editing techniques would fulfill all these criteria.

Targeted gene editing using CRISPR/Cas9 facilitates the introduction of donor DNA at specific loci in the genome, effectively tagging endogenous proteins of interest (14, 15). For neuronal cells, several CRISPR/Cas9-based knock-in strategies have been developed, relying on different mechanisms to repair double-stranded breaks (DSBs) introduced by Cas9. One strategy is based on homology-directed repair (HDR) to insert donor DNA into the genomic locus (16, 17). While this fatefully labels proteins of interest, HDR preferentially occurs during the S/G2 phases of the cell cycle, and is significantly downregulated in post-mitotic cells (18). This limits the application of HDR in post-mitotic neurons, although successful integration can still be observed with highly elevated donor DNA levels or via a combination of donor cleavage and micro-homology arms (19, 20). Additionally, in order to be efficient, HDR requires the addition of long homology arms to the donor DNA, which can be laborious to generate, and considerably complicates the development of knock-in constructs.

Alternative strategies are based on non-homologous end-joining (NHEJ) to repair DSBs, which is active throughout the cell cycle as well as in post-mitotic cells, and can be used to insert donor DNA with high accuracy (21–23). Based on NHEJ, the homology-independent targeted integration (HITI) method for endogenous protein tagging in post-mitotic neurons was previously developed and outperformed HDR-based methods in terms of efficiency and accuracy (20, 23). We hypothesized that this method would provide an accessible and scalable approach for the tagging of endogenous proteins in neurons. However, applications of this method have so far been limited to a few target proteins (20, 23, 24). In addition, designing and cloning of suitable knock-in constructs, compatibility of DNA delivery methods and validation have until now been quite challenging, and have not been addressed systematically.

Here, based on NHEJ we developed ORANGE, an Open Resource for the Application of Neuronal Genome Editing, that offers researchers the means to endogenously tag proteins of interest in post-mitotic neurons, allowing for the accurate investigation of protein expression, localization and dynamics. This toolbox includes 1) a single template vector that contains the complete knock-in cassette, which can be adapted in two straightforward cloning steps to tag virtually any protein of interest, and 2) a library of readily usable knock-in constructs targeting a set of more than 30 proteins. This library encompasses a wide variety of proteins, including cytoskeletal components, signaling molecules, endosomal markers, pre- and postsynaptic scaffolds, adhesion complexes, and receptors. We show that this tagging strategy facilitates single-cell labeling of synaptic proteins without overexpression artifacts and is compatible with multiple DNA delivery methods. Moreover, we demonstrate that this toolbox facilitates live-cell super-resolution imaging to accurately resolve endogenous protein localization and dynamics in neurons at high spatial and temporal resolution. Altogether, we present a robust, easy to implement toolbox for the tagging and visualization of endogenous proteins in post-mitotic neurons, allowing the in-depth investigation of diverse neuronal cell biological processes.

## RESULTS

### Generation of a CRISPR/Cas9-based knock-in toolbox for fluorescent tagging of endogenous proteins in neurons

In order to efficiently tag endogenous neuronal proteins, we first aimed to design a simple workflow to facilitate the rapid generation of knock-in constructs using conventional molecular cloning approaches. To this end, we designed a single CRISPR/Cas9 knock-in template vector (pORANGE), based on the original NHEJ-mediated HITI method (23). Our design allows the flexible insertion of a unique 20-nucleotide target sequence that guides Cas9 to the genomic locus of interest, and a donor sequence containing the knock-in sequence (e.g. GFP) to be inserted in the genomic locus (Figure 1A and S1). The generated knock-in construct contains all elements required for targeted CRISPR/Cas9-based genome editing: (1) a U6-driven expression cassette for the guide RNA (gRNA) targeting the genomic locus of interest, (2) the donor sequence containing the (fluorescent) tag, and (3) a Cas9 expression cassette driven by a universal β-actin promoter (Figure 1A). The donor sequence is generated by standard PCR, with primers that allow the introduction of a short linker and Cas9 target sequences flanking the donor (Figure 1A and S1A). These target sequences are identical to the genomic target sequence so that the same gRNA used to create a genomic DSB also mediates the removal of the donor DNA from the plasmid for genomic integration. Importantly, the orientation of the target sequence and PAM sites flanking the donor is inverted compared to the genomic sequence, to guarantee that integration occurs in the correct orientation (Figure S1A). For a detailed description of genomic target sequence selection, gRNA sequence and donor PCR primer design, we refer to the design and cloning protocol in the methods section (also see Figure S1). We found that this cloning strategy is easy to employ and allowed for the rapid and flexible generation of knock-in constructs.

**Figure 1.**
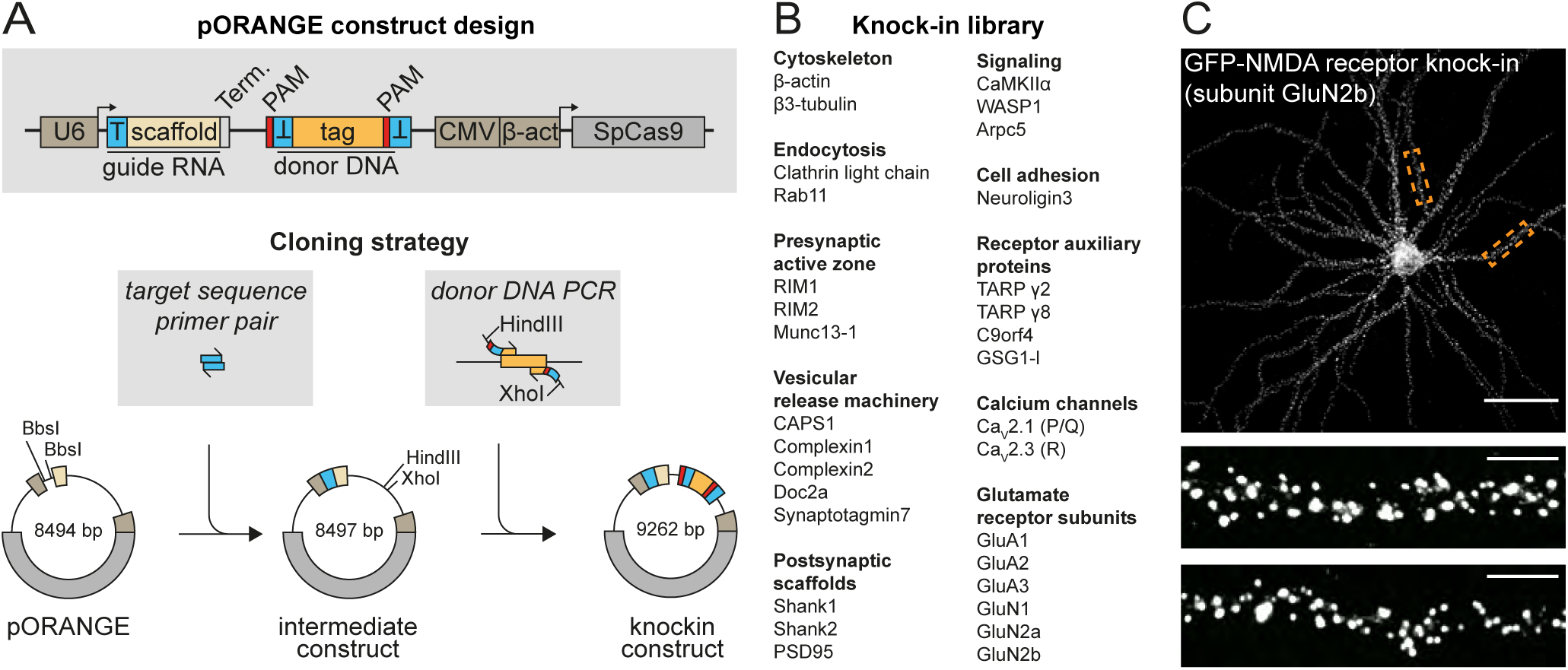
ORANGE: an easy to implement toolbox for endogenous tagging of proteins in neurons. **(A)** Overview of the pORANGE knock-in construct. T, indicates target sequence. Term., indicates termination sequence. First, the target sequence targeting the genomic locus of interest is ligated in the guide RNA cassette. Then, the donor sequence containing the tag of interest is inserted in the donor DNA cassette. **(B)** List of knock-in constructs generated based on the pORANGE template vector. **(C)** Example GFP-GluN2b knock-in neuron. Neurons were transfected at DIV 3 and imaged at DIV 21. Orange dashed boxes indicate zooms. Scale bars are 40 µm and 5 µm for the overview and zooms respectively.

Using the pORANGE template vector, we designed and generated a library providing more than 30 knock-in constructs to label endogenous neuronal proteins for fluorescence imaging (Figure 1B, C and 2). Throughout this study, when naming GFP knock-in constructs, we refer to the name of the protein that is labeled, where in N-terminally tagged proteins, GFP is in front of the protein name, and in C-terminally tagged proteins, the GFP is behind. To cover the many areas of neuronal cell biology, we selected proteins representing various molecular processes, including cytoskeletal components, intracellular signaling molecules and trafficking proteins (Figure 1B, 2). Furthermore, we put particular emphasis on synaptic components, and the library includes numerous endo- and exocytic proteins, pre- and postsynaptic scaffolds, cell adhesion complexes, calcium channels, receptor auxiliary proteins, and glutamate receptors. Together, our ORANGE toolbox includes a broad library of knock-in constructs and provides researchers an efficient strategy to adapt or design new constructs with relative ease, using standardized cloning techniques.

**Figure 2.**
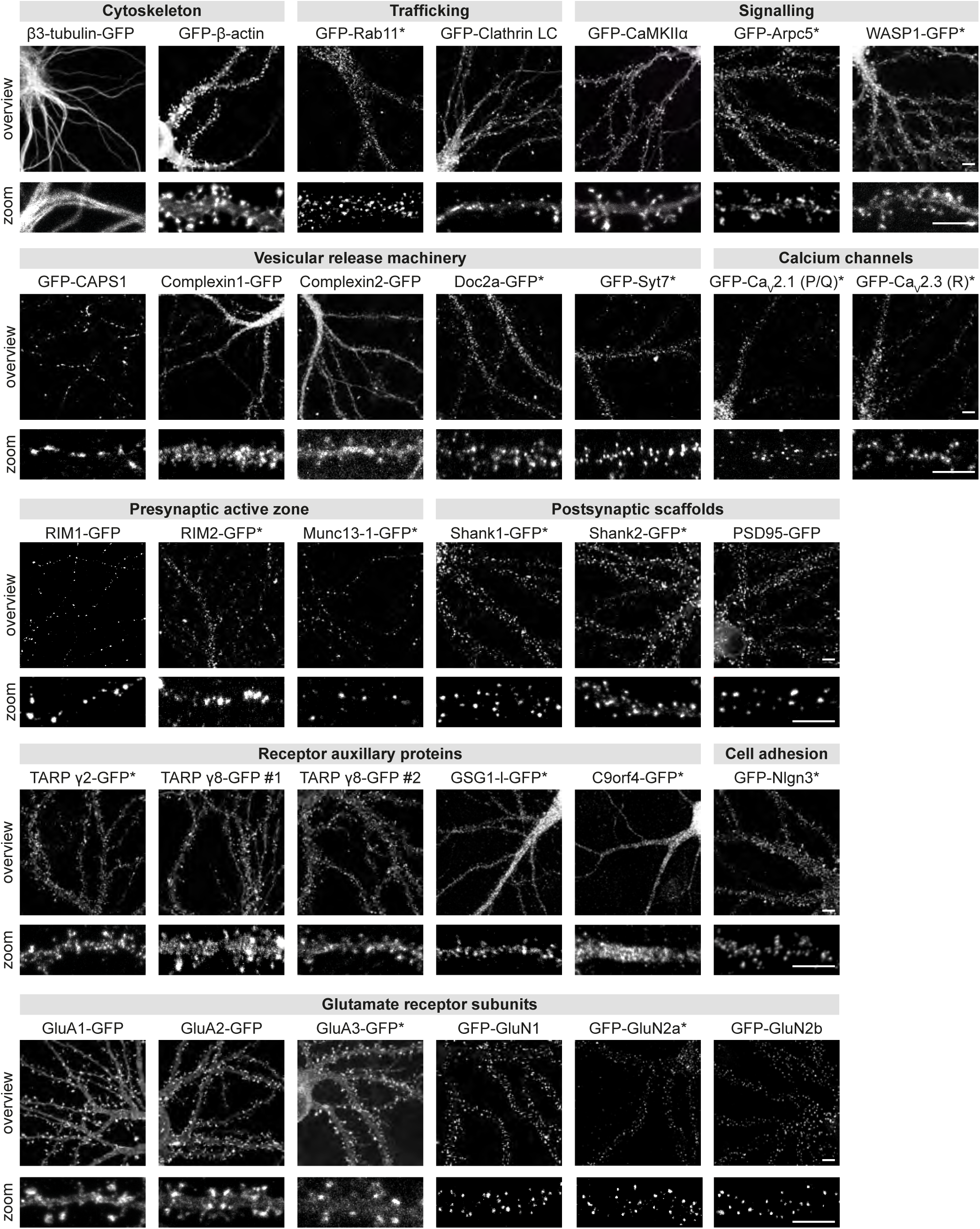
A versatile knock-in library for endogenous tagging of proteins in neurons. Representative images of CRISPR/Cas9 knock-ins in primary hippocampal rat neurons, categorized according to protein function or subcellular localization. Neurons were transfected at DIV 3 and imaged at DIV 21. *, indicates enhancement with anti-GFP antibody. Scale bars, 5 µm.

### ORANGE enables fast and efficient labeling of proteins at endogenous expression levels

To test the rate of donor integration and subsequent expression of tagged proteins, we co-transfected dissociated hippocampal neurons at day in vitro (DIV) 3 with a GFP knock-in construct for β3-tubulin (β3-tubulin-GFP knock-in) and a construct for soluble mCherry expression, making use of common lipofection-based DNA delivery methods (Figure S2). Because of the high protein turnover rate of β3-tubulin, integration of the donor should be rapidly observable by expression of the tagged protein. Successful labeling of β3-tubulin was observed within 24 hours after transfection, but at relatively low efficiency (1.1 ± 0.2% β3-tubulin GFP^+^/mCherry^+^ double positive cells). Labeling efficiency increased 10-fold over time and reached a plateau around 96 hours after transfection (10.9 ± 0.1% β3-tubulin GFP^+^/mCherry^+^; Figure S2), indicating that donor integration preferentially takes place within the first days after transfection.

Next, we determined the accuracy of our knock-in library. For none of the knock-in constructs in our library, we observed aberrant or diffuse expression of the integrated tag, indicating that off-target integration, or unintended GFP expression directly from the knock-in plasmid did not occur or is extremely rare. To get a general sense of the precision of donor integration into the targeted genomic locus, we determined the genomic sequence after integration for several knock-in constructs.

To express the knock-in vector in a large population of cells, we electroporated DIV0 neurons in suspension. We isolated genomic DNA from neuronal cultures 4 days after electroporation, and amplified the target locus with PCR primers around the 5’ and 3’ junctions of the site of integration. PCR products were ligated into pJET using blunt-end cloning, and individual clones were analyzed using Sanger sequencing (Figure S3). We were able to detect in-frame integration of the fluorescent tag in the targeted locus of almost all knock-ins that we evaluated, further demonstrating that the observed GFP signal likely originates from the intended GFP-fusion protein. Besides correct integration, we found various insertions and deletions leading to frameshift mutations. We noted that the frequency of indels was highly variable between different knock-ins, which is likely because the accuracy of Cas9 and NHEJ is reported to highly depend on the target sequence (25, 26). Given that we did not observe targeted neurons with non-specific GFP signal, or unexpected distribution of targeted proteins, these results indicate that while incorrect integration of the donor sequence can occur, this does not result in non-specific expression of the donor sequence. Thus, these analyses indicate that when observed, the GFP signal faithfully reports the distribution of the target protein.

To further determine whether the integrated fluorescent tag reliably labels the endogenous target protein, we chose to analyze the PSD95 knock-in construct in detail. PSD95 is a core postsynaptic scaffold molecule (27), for which specific antibodies are available. To edit the genomic PSD95 (*Dlg4*) locus before it is expressed, we transfected neurons with the PSD95-GFP knock-in construct well before synaptogenesis (DIV 3) and fixed at a mature stage (DIV 21) (Figure 3). In all neurons with detectable GFP signal, the GFP signal was found in a punctate pattern enriched in dendritic spines, characteristic for endogenous PSD95 expression. The GFP signal closely co-localized with anti-PSD95 antibody staining, resulting in a strong correlation between intensity of PSD95 immunostaining and GFP intensity in PSD95-GFP knock-in neurons (Pearson r: 0.72 R^2^: 0.51 *P* < 0.001, n = 450 synapses from 9 neurons; Figure 3B), indicating that the observed GFP signal reliably reports the localization and labels the complete protein pool of PSD95. To test if the knock-in affects total PSD95 levels, we used the PSD95 antibody staining to compare protein levels between PSD95-GFP knock-in and control neurons that were transfected with soluble GFP (GFP control). Although we observed that in a subpopulation of PSD-95 knock-in neurons protein levels were modestly lower, in almost all neurons, PSD95 levels in PSD95-GFP knock-in neurons (relative fluorescence intensity: 0.85 ± 0.04, n = 17 neurons) were comparable to GFP control (0.98 ± 0.02, n = 15 neurons, ANOVA, *P* > 0.05) (Figure 3C; inset). In contrast, overexpression of PSD95-GFP led to significantly increased, and more variable, synaptic PSD95 protein levels (relative fluorescence intensity: 4.2 ± 0.4, n = 17 neurons, *P* < 0.001). Moreover, synapse size was significantly increased in neurons overexpressing PSD95 (0.18 ± 0.01 µm^2^) compared to GFP control (0.13 ± 0.01 µm^2^, ANOVA, *P* < 0.001), but was unaffected in PSD95-GFP knock-in neurons (0.14 ± 0.001 µm^2^, *P* > 0.05; Figure 3D). Lastly, we found that PSD95 was significantly more enriched at synapses in PSD95 knock-in cells (ratio synapse/shaft intensity: 17.6 ± 1.6) compared to PSD95 overexpressing neurons (11.8 ± 1.0, Student’s t-test, *P* < 0.01; Figure 3E), indicating that a large fraction of overexpressed PSD95 mis-localized to the dendritic shaft.

**Figure 3.**
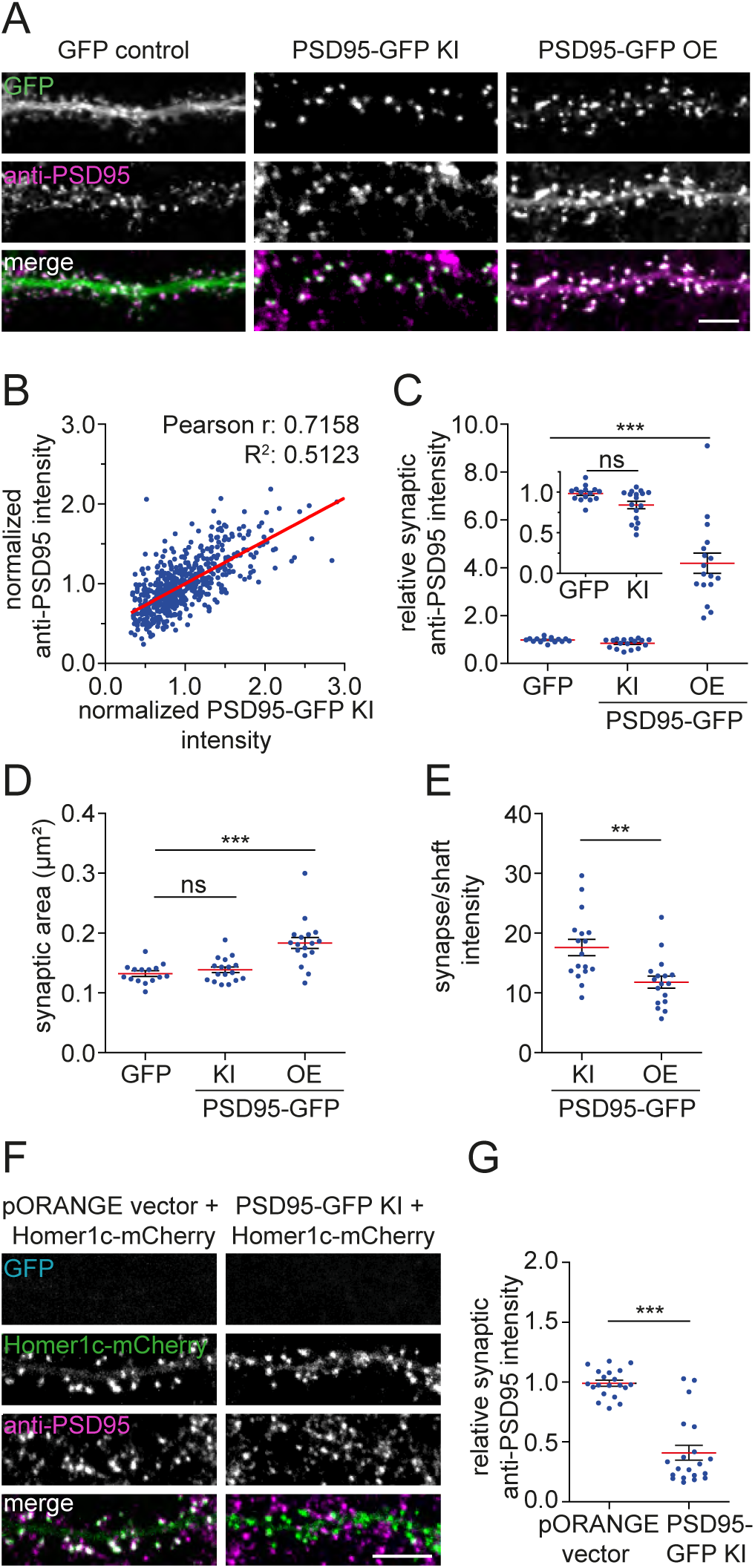
Validation of CRISPR/Cas9 targeting efficiency. **(A)** Representative images of dendrites transfected with soluble GFP, PSD95-GFP knock-in (KI) construct, or a PSD95-GFP overexpression construct (green) stained with anti-PSD95 (magenta). DIV 21. Scale bar, 5 µm. **(B)** Correlation between PSD95-GFP knock-in and anti-PSD95 staining intensity. **(C)** Quantification of synaptic PSD95 levels, **(D)** synapse area, and **(E)** PSD95 synapse/shaft intensity. **(F)** Representative images of dendrites co-expressing Homer1c-mCherry (green) and either the empty pORANGE template vector or PSD95-GFP knock-in construct (blue) stained with anti-PSD95 (magenta). DIV 21. **(G)** Quantification of PSD95 levels in transfected, but knock-in negative neurons. Scale bar, 5 µm. Data are represented as mean ± SEM. ns, indicates not significant *, indicates P < 0.05, **, indicates P < 0.01, ***, indicates P < 0.001, ANOVA or t-test.

Transfection of the knock-in construct did not always result in successful knock-in of GFP (Figure S2, S3). To determine whether in transfected, but GFP knock-in-negative neurons, integration of the GFP tag was simply not successful, or whether integration had introduced mutations affecting protein expression, we co-transfected neurons with the PSD95-GFP knock-in construct and a Homer1c-mCherry overexpression construct. Next, we measured PSD95 levels in Homer1c-mCherry positive neurons that did not show detectable PSD95-GFP signal (Figure 3F, G). In most of these neurons, PSD95 protein levels were significantly downregulated (relative fluorescence intensity: 0.41 ± 0.06, n = 20 neurons), compared to co-transfection of the empty pORANGE template vector together with Homer1c-mCherry (0.99 ± 0.02, n = 20 neurons, Student’s t-test, *P* < 0.001) suggesting partial or complete knockout of the target protein in these neurons, probably due to frame-shift mutations or truncations. Thus, while successful knock-in results in accurate detection of endogenous PSD95, erroneous integration may lead to partial loss-of-function of the targeted gene in knock-in negative neurons. Altogether, these data demonstrate that ORANGE enables successful integration of fluorescent tags at the targeted genomic locus, resulting in expression of fusion proteins which reliably reports the localization of proteins of interest at endogenous levels.

### Lentiviral mediated delivery of knock-in constructs enables labeling of endogenous proteins in organotypic hippocampal slice cultures

We achieved successful genomic integration of donor sequence in primary neuronal cultures using standard transfection methods (lipofection or electroporation), while previous studies mainly used adeno-associated virus (AAV)-based approaches (19, 22, 23). However, it remained unclear whether lentivirus-mediated infection of neurons would deliver sufficient copy numbers of the donor sequence for efficient genomic integration in primary neuronal cultures and organotypic hippocampal slices. To address this, we divided the ORANGE knock-in cassette over two lentiviral constructs (Figure 4A), because the full cassette exceeds the packaging limit of lentiviral particles (LVs). Also, premature co-expression of Cas9 and the gRNA during the production of viral particles in packaging cells would lead to removal of the donor DNA. We infected dissociated hippocampal cultures with Cas9 and knock-in LVs targeting the genes encoding the AMPA receptor subunit GluA1 and β3-tubulin. GluA1-GFP and β3-tubulin-GFP positive neurons were observed in hippocampal cultures after double infection with knock-in and Cas9 LVs (Figure 4B), demonstrating that this lentiviral approach can mediate successful genomic integration. We next injected the CA1 region of mouse organotypic hippocampal slice cultures with knock-in LVs for GluA1-GFP at DIV 1 and analyzed GFP fluorescence at DIV 10. We observed a number of GluA1-GFP positive CA1-pyramidal neurons with GFP signal largely restricted to dendritic spines, with weaker signal on dendritic shafts as well as the soma (Figure 4C,D), consistent with the synaptic enrichment of GluA1-containing AMPA receptors in excitatory synapses. Based on location of the cell body and dendritic morphology, we also identified interneurons that were positive for GluA1-GFP. In these neurons, that generally lack dendritic spines, punctate GluA1-GFP labeling was restricted to dendritic shafts (Figure 4D). Thus, LVs can be used to label endogenous proteins in both cultured and organotypic neuronal preparations, broadening the potential applications of this CRISPR/Cas9 genome editing toolbox.

**Figure 4.**
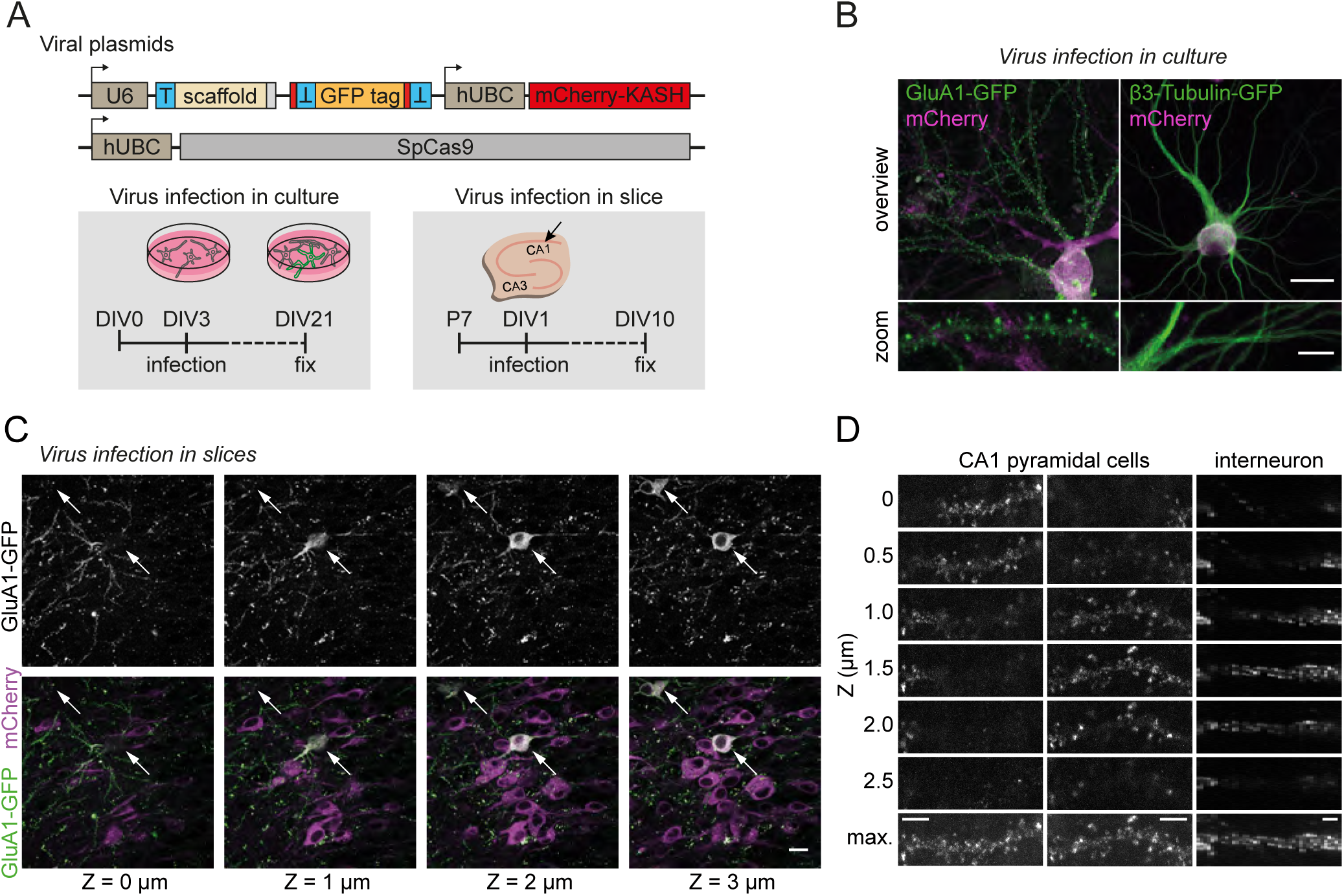
ORANGE knock-in in culture and organotypic slices using dual-lentiviral approach. **(A)** Overview of lentiviral constructs and timeline showing age of infection and fixation. **(B)** Representative images of infected (magenta) primary rat hippocampal neurons positive for GluA1-GFP knock-in or β-tubulin knock-in (green). Scale bars, 20 µm and 5 µm for the overview and zooms respectively. **(C)** Representative images of GluA1-GFP knock-in in organotypic hippocampal slices from mice. Shown are a series of individual 1 µm planes from a Z-stack. Arrows indicate GFP positive cells. Scale bar, 20 µm. **(D)** Representative zooms of GluA1-GFP knock-in dendrites from a CA1 pyramidal cell and an aspiny interneuron. Shown are individual 0.5 µm planes from a Z-stack and the maximum-projection (max). Scale bar, 2 µm.

### ORANGE-mediated knock-in allows super-resolution imaging of endogenously expressed proteins in neurons

We envisioned that tagging endogenous proteins with fluorophores particularly presents advantages for super-resolution imaging by facilitating labeling of proteins in a subset of neurons, and overcoming many artefacts associated with immunolabeling or overexpression of tagged proteins. To evaluate this, we employed our PSD95-GFP and GFP-β-actin knock-in constructs to resolve and correlate their subcellular organization in individual neurons using gated stimulated-emission depletion (gSTED) microscopy.

First, we performed two-color gSTED on PSD95-GFP knock-in neurons stained with anti-PSD95 antibodies to assess the degree of co-localization (Figure 5A). Both the confocal and STED images of individual synapses revealed a high degree of co-localization between the PSD95-GFP knock-in and anti-PSD95 staining (Figure 5A-E). Additionally, gSTED revealed that even at the subsynaptic level, PSD95-GFP knock-in and PSD95 immunolabeling co-localized. Second, we assessed the degree of co-localization between the GFP-β-actin knock-in and anti-PSD95 staining. We used phalloidin staining to determine whether F-actin composition of spines was unaltered in GFP-β-actin knock-in neurons (Figure S4A-C). Indeed, GFP-β-actin knock-in signal intensity strongly correlated with phalloidin intensity (Pearson r: 0.73, R^2^: 0.53, *P* < 0.001, n = 450 spines from 9 neurons; Figure S4B), and we did not find differences in average phalloidin staining intensity between control (relative fluorescence intensity 1.02 ± 0.03, n = 9 neurons) and knock-in neurons (1.04 ± 0.02, n = 9 neurons, ANOVA, *P* > 0.05) (Figure S4C), indicating that the GFP-β-actin knock-in accurately labels endogenous actin. While not visible in the confocal images, two-color gSTED of GFP-β-actin knock-in, co-stained with anti-PSD95 staining as synaptic marker revealed that although β-actin is enriched in dendritic spines, it is largely excluded from the PSD (Figure 5F-J). To analyze the degree of co-localization we used two independent metrics: the Pearson’s correlation coefficient (PCC, Figure 5K) and Manders’ overlap correlation (MOC) (28, 29). For the MOC, two different values are derived, *M1;* the fraction of knock-in overlapping anti-PSD95 staining (Figure 5L), and *M2;* the fraction of anti-PSD95 staining overlapping the knock-in (Figure 5M). The PCC and MOC of the PSD95-GFP knock-in and anti-PSD95 staining, as measured with confocal microscopy, revealed a high degree of co-localization (confocal: median PCC = 0.95, *M1* = 0.75, *M2* = 0.80, n = 10 neurons; Figure 5K-M). Because gSTED microscopy resolved the subsynaptic distribution at higher resolution, these values were lower (STED: median PCC = 0.88, ANOVA, *P* < 0.001, *M1* = 0.63, *P* < 0.001, *M2* = 0.70, *P* >0.05; Figure 5K-M). Using STED microscopy the distributions of β-actin and PSD95 revealed a non-uniform distribution of β-actin that surrounded, rather than co-localized with PSD95 as appeared from the confocal images, with consequent lower PCC and MOC values (n = 7 neurons, STED: median PCC = 0.78, ANOVA, *P* < 0.001, *M1* = 0.19, *P* < 0.001, *M2* = 0.47, *P* < 0.001; confocal: median PCC = 0.88, *P* < 0.001,, *M1* = 0.50, *P* < 0.001, *M2* = 0.77, *P* >0.05) (Figure 5K-M).

**Figure 5.**
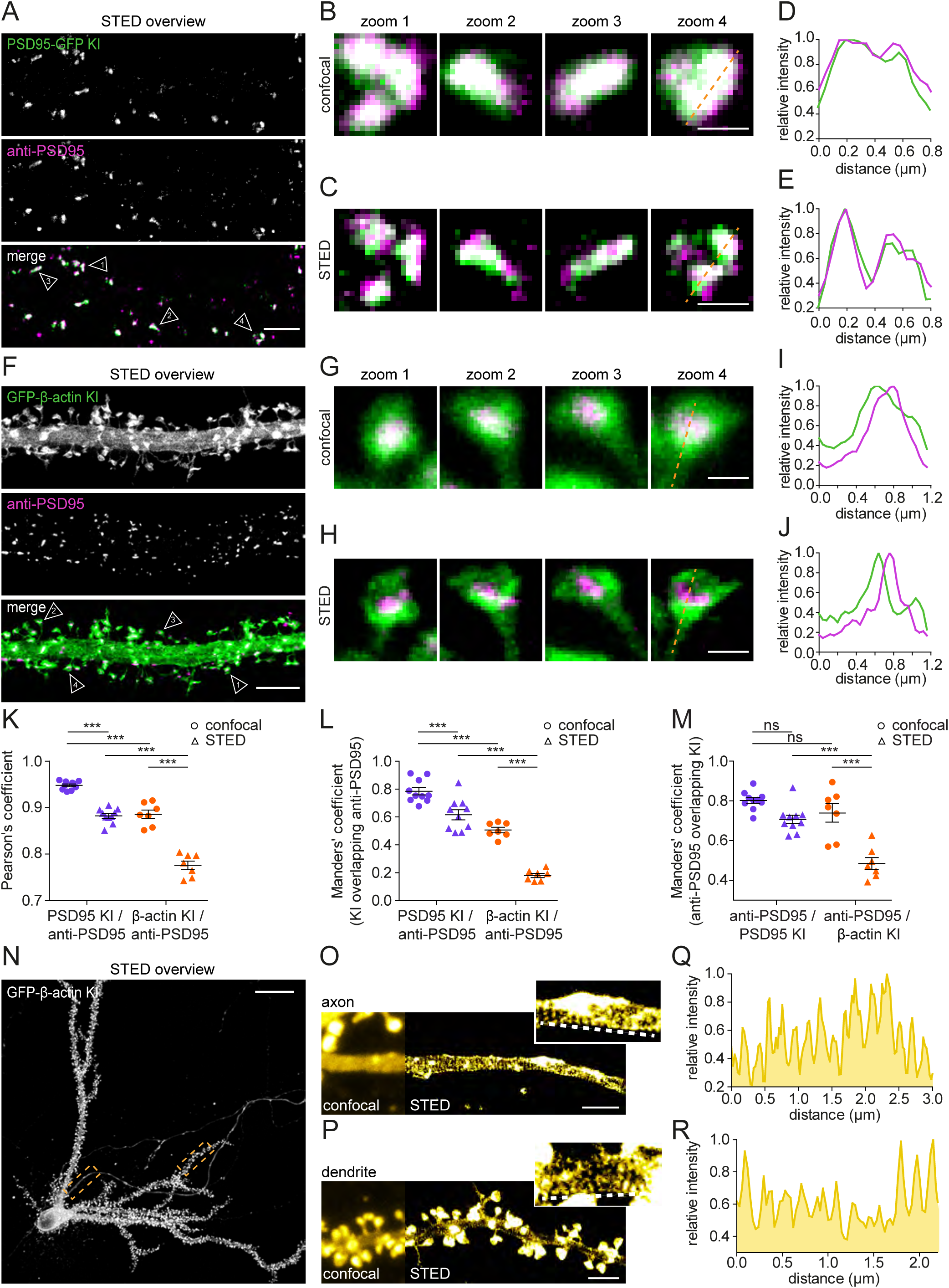
Super-resolution microscopy investigation of the subcellular distribution of endogenous proteins in single neurons. **(A)** Representative gSTED images of dendrites positive for PSD95-GFP knock-in enhanced with anti-GFP (ATTO647N; green) and anti-PSD95 (Alexa594; magenta). DIV 21. Scale bar, 2 µm. **(B, C)** Zooms from (A) of individual synapses resolved with confocal (B) and gSTED (C). Scale bar, 500 nm. **(D, E)** Line scans of confocal (D) and gSTED (E) images shown in (B) and (C) respectively. **(F)** Representative gSTED images of dendrites positive for GFP-β-actin knock-in stained with anti-GFP (ATTO647N; green) and anti-PSD95 (Alexa594; magenta). DIV 21. Scale bar, 2 µm. **(G, H)** Zooms from (F) of individual spines resolved with confocal (G) and gSTED (H). Scale bar, 500 nm. **(I, J)** Line scans of confocal (I) and gSTED (J) images shown in (G) and (H) respectively. **(K)** Pearson’s correlation coefficient quantifying co-localization between PSD95-GFP or GFP-β-actin knock-in with anti-PSD95 staining. **(L,M)** Manders’ correlation of PSD95-GFP or GFP-β-actin knock-in overlapping with PSD95 staining (L), or anti-PSD95 staining overlapping with PSD95-GFP or GFP-β-actin knock-in (M). **(N)** Representative gSTED image of a GFP-β-actin knock-in neuron enhanced with anti-GFP (ATTO647N). Cytosolic monomeric β-actin was removed using an extraction protocol. DIV 21. Scale bar, 20 µm. **(O, P)** Zooms of axon and dendrites as indicated with boxes in (N), comparing confocal and gSTED imaging. Scale bar, 2 µm. **(Q, R)** Line scans from zooms in (O) and (P) respectively. Data are represented as mean ± SEM. ns, indicates not significant, ***, indicates P < 0.001, ANOVA.

Recent super-resolution studies have demonstrated that the F-actin cytoskeleton forms ring-like structures that are periodically organized along axons, as well as dendrites (30–32). Typically, in experiments investigating these structures, F-actin is visualized by phalloidin staining, which does not permit single-cell labeling. Therefore, neurons are often grown in low-density cultures, which has profound effects on neuronal development (33, 34). Thus, to explore the advantages of single-cell labeling using ORANGE even further, we tested whether we could resolve this particular organization of the actin cytoskeleton in individual β-actin knock-in neurons within dense, mature neuronal cultures. Using gSTED imaging we observed distinct periodic actin structures in both the axon and dendrites (Figure 5N-R). Furthermore, using two-color gSTED imaging on β3-tubulin-GFP knock-in neurons immunolabeled with anti-α-tubulin allowed us to resolve the neuronal microtubule network (Figure S4D-F). Altogether, these results demonstrate that ORANGE allows for direct visualization of nanoscale macromolecular structures at conditions of endogenous expression and importantly, in individual labeled neurons.

### Live-cell single-molecule tracking of endogenous intracellular proteins

In addition to imaging fixed cells, recombinant fluorescent tags provide a powerful approach to investigate endogenous protein dynamics within living cells. To demonstrate this directly, we designed knock-in constructs targeting the gene encoding CaMKIIα, an abundant neuronal Ca^2+^-activated signaling protein essential for learning and memory (35), to map the nanoscale distribution and dynamics of CaMKIIα in live neurons. First, we made a GFP knock-in construct tagging CaMKIIα, and consistent with previous studies (17, 36), confocal and two-color gSTED microscopy showed that the GFP signal was primarily cytoplasmic with moderate enrichment in spines (Figure 2 and S4G-J). We next replaced GFP for mEos3.2, a photoconvertible protein compatible with single-molecule tracking based on photoactivated localization microscopy (PALM) (37, 38) (Figure S5). Individual mEos3.2-CaMKIIα molecules were tracked to reconstruct a super-resolved image of CaMKIIα distribution (Figure S5B) and to map single-molecule trajectories in spines and dendrites (Figure S5C, D). From this, we calculated the mean-squared displacements to derive the diffusion coefficient for individual trajectories (Figure S5E, F), revealing two distinct dynamic CaMKIIα populations: a mobile population (mean diffusion coefficient 0.145 ± 0.049 µm^2^/s) and a slower moving population (0.0140 ± 0.0011 µm^2^/s, n = 11 neurons). Thus, genetic tagging with photoconvertible molecules such as mEos3.2 allows live-cell single-molecule tracking PALM experiments to map the distribution and dynamics of endogenous, intracellular neuronal proteins.

### Nanometer dissection of endogenous NMDA receptor distribution and dynamics within individual synapses

Given that overexpression of synaptic proteins can have detrimental consequences for synapse morphology and function, we next tested whether we can genetically label endogenous synaptic receptors for live-cell super-resolution imaging. Overexpression of individual receptor subunits significantly affects subunit stoichiometry of endogenous receptors, leading to alterations in their trafficking and ultimately the physiology of synapses (39). In particular, since the local density and subsynaptic positioning of glutamate receptors are key determinants of synaptic transmission and plasticity, it is critical to study the positioning of these receptors at endogenous expression levels (1, 40). Based on overexpression and antibody-labeling studies, the spatial organization of NMDA receptors at excitatory synapses has been proposed to be heterogenous, with receptors accumulating in distinct subsynaptic nanodomains (41–43). Therefore, we aimed to combine ORANGE with super-resolution techniques to dissect the distribution of dynamics of NMDA receptors, which are critical for the induction of synaptic plasticity underlying development, learning, and memory formation. NMDA receptors are heteromeric protein complexes, with varying subunit composition, but contain at least two copies of the GluN1 subunit. Thus, to tag the total pool of NMDARs, we developed a knock-in construct to tag endogenous GluN1 subunits at the N-terminus with GFP (Figure 6A). Several studies have consistently estimated that the number of NMDA receptors at individual synapses is relatively low, ranging from 10 to 20 receptor complexes per synapse (27, 44). Despite these low copy numbers, we could detect dendritic clusters of GFP-GluN1, most of which colocalized with immuno-labeled PSD95 (Figure 6B, C). Interestingly, we found that GFP-GluN1 intensity did not correlate with anti-PSD95 immunolabeling intensity (Figure 6D) (Pearson r: 0.19, R^2^: 0.038, n = 450 GluN1 clusters from 9 neurons), consistent with earlier studies that showed that the total number of NMDA receptors is largely invariable and does not scale with synapse size (45–47). Using gSTED imaging, we found that while most GFP-GluN1 clusters localized to synapses, some smaller extrasynaptic clusters could be detected (Figure 6E-G). Next, we measured the total GFP-GluN1 cluster area in individual synapses, and found a slight correlation with synapse size (Pearson r: 0.64, R^2^: 0.4087, n = 266 synapses from 3 neurons; Figure 6H). Thus, our data suggest that the subsynaptic area covered by NMDARs, but not the total number of receptors, scales with synaptic size. gSTED imaging of individual synapses also indicated that the subsynaptic distribution of GFP-GluN1 is heterogeneous (Figure 6F, I), with individual synapses containing one or more smaller GFP-GluN1 substructures (Figure 6J) (n = 266 synapses from 3 neurons).

**Figure 6.**
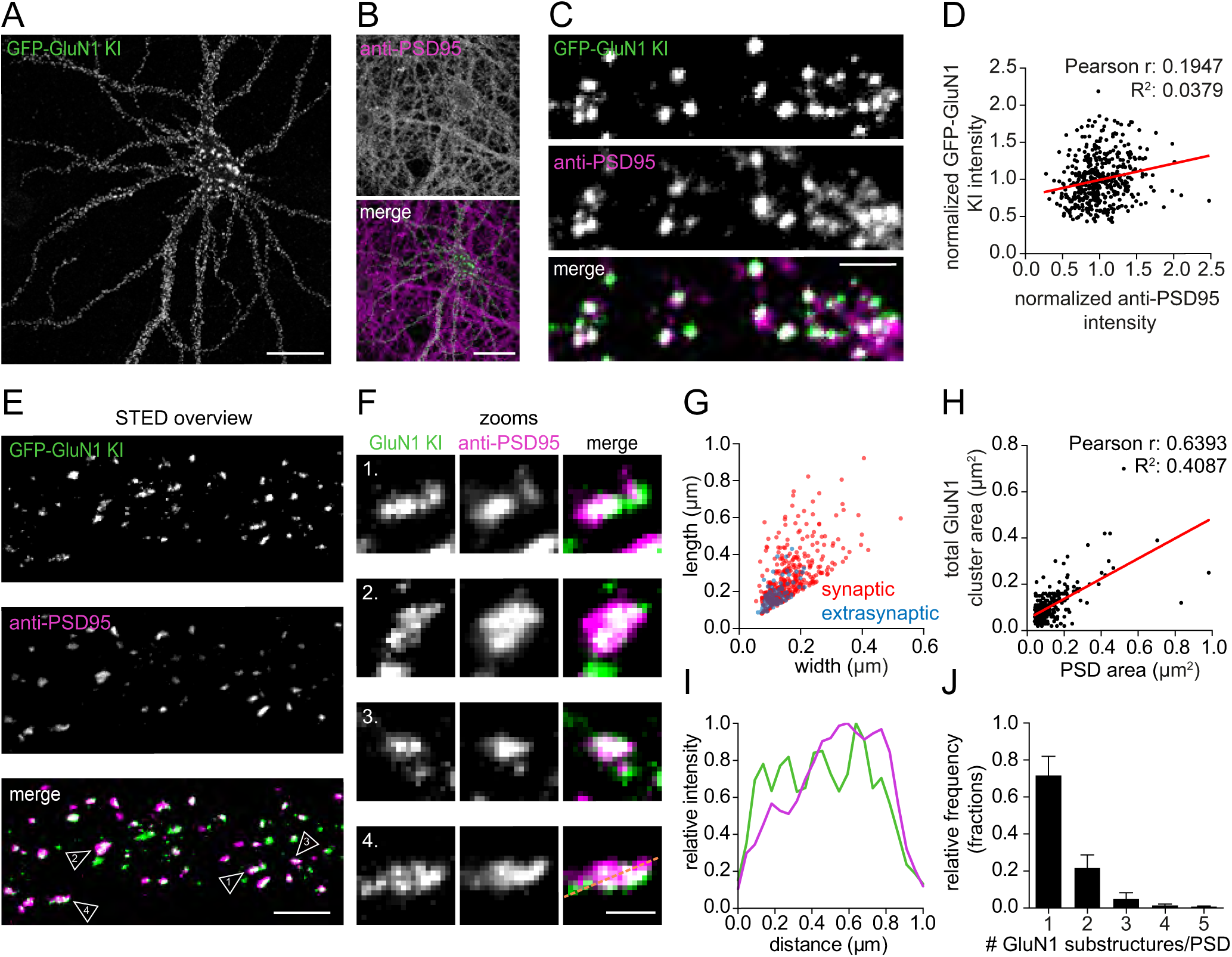
Resolving the sub-synaptic distribution of endogenously tagged NMDA receptors. **(A, B)** Representative confocal image of a GFP-GluN1 knock-in (green) neuron, and stained for endogenous PSD95 (magenta) (B). DIV 21. Scale bars in (A) and (B), 10 µm. **(C)** Representative images of dendrites positive for GFP-GluN1 knock-in (ATTO647N; green) stained for PSD95 (Alexa594; magenta). Scale bar, 2 µm. **(D)** Correlation between GFP-GluN1 knock-in and anti-PSD95 staining intensity within GFP-GluN1 puncta. **(E)** Representative gSTED images of dendrites positive for GFP-GluN1 knock-in enhanced with anti-GFP (green), and anti-PSD95 (magenta). DIV 21. Scale bar, 2 µm. **(F)** Zooms of individual synapses indicated in (E). Scale bar, 500 nm. **(G)** Full width at half maximum (FWHM) analysis of GFP-GluN1 structures comparing width and length of individual synaptic (red) and extrasynaptic (blue) GluN1 clusters. **(H)** Correlation between GFP-GluN1 cluster area and synapse area (based on anti-PSD95 staining) for individual synapses. **(I)** Line scan of synapse zoom 4 in (F). **(J)** Quantification of the GFP-GluN1 substructures per synapse. Data are represented as mean ± SEM.

To further investigate the subsynaptic distribution of NMDA receptors, we turned to single-molecule localization microscopy (SMLM). GFP-tagged GluN1 was immunolabeled with anti-GFP and Alexa-647-coupled secondary antibodies and imaged using direct stochastic optical reconstruction microscopy (dSTORM) to reconstruct the spatial organization of NMDA receptors at individual synapses with nanometer resolution (Figure 7A, B). Clusters of GFP-GluN1 receptors were identified using DBSCAN (48). Next, all localizations within individual clusters were plotted, and color-coded for the local density. These local density maps revealed that within individual clusters NMDA receptors form distinct nanodomains (Figure 7C), consistent with our gSTED data showing a heterogeneous distribution within the PSD (Figure 6F-J). Quantitative analysis of these nanodomains demonstrated that the majority of GFP-GluN1 clusters contained one to three nanodomains with a median size of ∼60 nm (IQR: 53 – 71 nm) (n = 859 GFP-GluN1 clusters from 3 neurons) (Figure 7D, E). Thus, these SMLM data indicate that endogenous NMDA receptors form distinct subsynaptic nanodomains.

**Figure 7.**
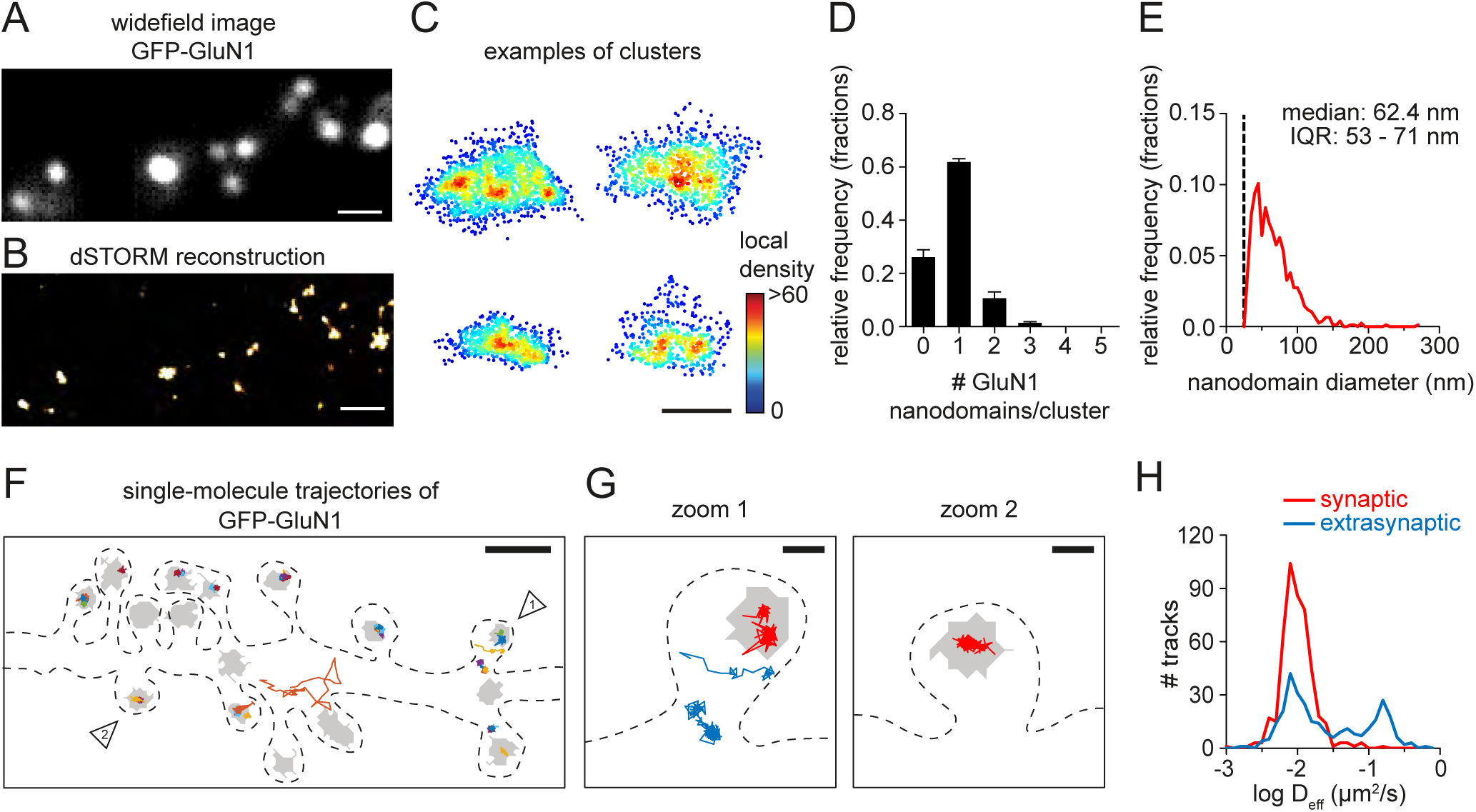
NMDA receptors cluster into nanodomains and are highly immobilized in synapses. **(A)** Representative image of dendrite positive for GFP-GluN1 knock-in enhanced with anti-GFP (Alexa647). DIV 21. Scale bar, 1 µm. **(B)** Single-molecule dSTORM reconstruction of example shown in (A). Scale bar, 1 µm. **(C)** Examples of individual GFP-GluN1 clusters, with single localizations plotted and color-coded based on the local density. Scale bar, 200 nm. **(D)** Quantification of number of GFP-GluN1 nanodomains per cluster. **(E)** Frequency distribution of GFP-GluN1 nanodomain size. Dotted line indicates nanodomain size cutoff. Bin size: 5 nm. **(F)** Representative example of GFP-GluN1 single-molecule trajectories in a dendrite plotted with a random color and on top of a thresholded PSD mask (grey) based on Homer1c-mCherry widefield image. Dotted line indicates cell outline. DIV 21. Scale bar, 1 µm. **(G)** Zooms of individual spines indicated in (F), with example trajectories of synaptic (red), or extrasynaptic (blue) receptors. Scale bar, 200 nm. **(H)** Frequency distribution showing the diffusion coefficient of synaptic and extrasynaptic tracks. Data are represented as mean ± SEM.

To gain insight in the subsynaptic mobility of endogenously expressed NMDA receptors, we probed the diffusion kinetics of individual receptors using live-cell single-molecule tracking or universal point accumulation in nanoscale topography (uPAINT) (49). Stochastic labeling of individual GFP-tagged receptors with a GFP-nanobody coupled to ATTO-647N, provided a map of individual receptor mobility along stretches of dendrites (Figure 7F,G). Consistent with our previous observations, we found most receptor trajectories within the boundaries of the PSD. Strikingly, we found that these synaptic NMDA receptors were largely immobilized (median diffusion coefficient synaptic tracks: 0.0096 µm^2^/s IQR: 0.0079-0.0122, n = 462 tracks from 6 neurons) while on average, extrasynaptic receptors diffuse at higher rates (0.0224 µm^2^/s IQR: 0.0123-0.0419, n = 307 tracks from 6 neurons). Altogether, by combining the ORANGE toolbox with super-resolution microscopy, we show that NMDA receptors are enriched in the PSD, where they are highly immobilized and cluster in subsynaptic nanodomains

## DISCUSSION

Mapping the subcellular distribution of proteins at high spatial resolution is fundamental to understand cell biological processes. Ongoing developments in super-resolution imaging technologies have dramatically improved the spatial resolution, allowing the dissection of molecular organization of subcellular structures at nanometer precision. A major obstacle remains the availability of a flexible strategy to efficiently and specifically label endogenous proteins, especially in neurons. Here, we developed ORANGE, a simple and scalable toolbox for epitope tagging of endogenous proteins using CRISPR/Cas9, and we provide a readily usable knock-in library that allows in-depth interrogation of protein distribution and dynamics in post-mitotic neurons at high spatial resolution. While CRISPR/Cas9 based tagging approaches have been developed for neurons, until now, large-scale applications and further development of these methods have been limited. ORANGE is easy to implement and scalable, only requiring standard cloning methods. Moreover, we have demonstrated that this approach is compatible with various generally used DNA delivery methods including lipofection, electroporation and LVs and thus, can be used in both primary neuronal cultures as well as organotypic hippocampal slices.

We demonstrated the level of accuracy of targeted genomic integration using ORANGE in several ways. First, for all our targets we found that the distribution of the GFP signal was consistent with previous reports of protein localization inferred from immunohistochemistry, or biochemical fractionation experiments. Second, for the well-characterized proteins PSD95 and β-actin, we found a high degree of co-localization and significant linear correlation between the knock-in GFP signal and endogenous staining intensity using antibodies and organic dyes. Even when resolved with super-resolution STED, we found that subsynaptic PSD95-GFP puncta overlapped closely with endogenous PSD95 staining. Third, we analyzed at the genomic level whether insertion of GFP was correct. Although the level of indels is highly variable between individual targets, we identified positive clones for most knock-ins. Thus, even though indels are generated in a subset of the transfected neurons, this should not form a limitation for many purposes, including fluorescent imaging as only neurons with detectable fluorescent signal are selected. Importantly, our results indicated that when expressed, the tag virtually always accurately reports protein localization and in most knock-in positive neurons, does not affects protein levels. We did observe that for a small number of cases protein levels in knock-ins were slightly lower compared to wildtype cells. This may indicate that, in these cells, one of the two alleles contains indels after genome editing, and/or failed to integrate the donor DNA, consistent with estimates that with the HITI method, 30-50% of knock-in positive cells show biallelic integration (23). Ongoing advancements in CRISPR/Cas9 technology are likely to lead to new developments that increase the on-target integration efficiency and precision of this approach. For instance, Cas9 variants that are more accurate could increase specificity and decrease indel frequency (50, 51), and the indel frequency may be predicted based on the target sequence (25, 26). Also, alternative delivery methods such as ribonucleoproteins (RNPs) (52), might increase the efficiency of targeted integration.

An important advantage of our method is that targeted integration of common epitope tags circumvents the need for developing new specific antibodies. In particular, for proteins that are highly homologous in their amino acid sequence, and for which generating specific antibodies is challenging, it is now possible to develop specific knock-in constructs that will report subcellular localization at unmatched specificity. As an example, we demonstrated successful knock-ins for RIM1 and RIM2, two highly homologous active zone proteins for which isoform-specific antibodies are not available. Thus, instead of relying on individual antibodies with varying levels of specificity and efficiency, the ORANGE toolbox presented here makes it possible to use universal antibodies recognizing epitope tags, or fluorescent proteins that directly report protein localization. Most of these tools are well-characterized and have been optimized for a number of applications, thereby circumventing cumbersome optimization procedures, and facilitating the comparison of results between different preparations, experimental setups and laboratories. The knock-in constructs presented in our library are designed using the rat genome as a template. But, due to high gene homology, multiple of the knock-in constructs, are compatible with the mouse genome and human (see Table S1). For example, we have shown that our GluA1-GFP knock-in works both in rat hippocampal cultures, as well as mouse organotypic hippocampal slices.

ORANGE is easily employed on targets that are yet uncharacterized. Deep sequencing efforts and high-resolution proteomics studies continue to implicate novel proteins in biological processes, but for many of these proteins specific and efficient antibodies are lacking. For instance, we developed knock-in constructs for two AMPA receptor complex constituents, C9orf4 and GSG1L, that have only recently been discovered in a high-resolution proteomics study (53). For both proteins functional characterization is available (54, 55), but high-resolution information on subcellular distribution is still lacking due to unavailability of efficient antibodies.

The ability to tag endogenous proteins in sparse subsets of cells is particularly advantageous for super-resolution approaches. The unmatched resolution of these approaches will detect any distortion in molecular organization due to for instance overexpression artefacts, and therefore these methods are highly sensitive to non-specific labeling. Also, sparse labeling of cells increases contrast and provides internal negative controls, as neighboring, non-targeted cells are unlabeled. We exploited these advantages to dissect the subcellular distribution of a number of neuronal proteins using different super-resolution imaging approaches. We mapped the distribution of endogenous cytoskeletal elements, signaling proteins and synaptic neurotransmitter receptors. Specifically, we found that endogenous actin is organized in a periodic pattern both in axons and dendrites, consistent with previous observations (30–32). Additionally, in live neurons we were able to track the dynamic behavior of the endogenous signaling protein CaMKII and NMDA-type glutamate receptors. Our experiments demonstrate that endogenous CaMKIIα has two distinct kinetic populations. Focusing on glutamate receptors, we found that endogenous NMDA receptors are highly immobilized at synaptic sites, and enriched in distinct subsynaptic nanodomains. This particular distribution is likely to shape the efficiency of receptor activation by presynaptically released neurotransmitters (1), and therefore dissection of the underlying molecular mechanisms is essential for the understanding of synapse physiology. Thus, ORANGE enables live-cell single-molecule tracking for virtually any protein, using photoconvertible fluorophores for PALM of intracellular proteins, and nanobody-based uPAINT to track surface receptors, and thus provides a scalable approach to efficiently and reliably map the dynamic distribution of endogenous proteins at nanometer resolution in live neurons.

We believe that ORANGE is a simple and efficient genome editing toolbox that will rapidly advance various fields in biology, by allowing the in-depth investigation of protein distribution in cultured cell lines, primary cells, organotypic slices, and animal models, but in particular, it presents one of the few possibilities to tag proteins in neurons. Further development of tools for *in vivo* gene editing combined with cell-type specific targeting of epitope tags would allow interrogation of protein distribution in specialized neuron types in the brain. Apart from epitope tagging, our toolbox can for example be used for insertion of enzymes for proximity biotinylation (56), labeling of organelles for electron microscopy (57) or light-sensitive dimerization sequences for optical control over protein or organelle positioning (58, 59). The unprecedented number of applications of ORANGE will undoubtedly deepen our molecular understanding of how the spatial distribution of endogenous proteins contributes to cell biological processes.

## ACKNOWLEDGMENTS

We want to thank Lisa A.E. Catsburg for contributing knock-in constructs to the library and all members of the MacGillavry lab for discussions. We want to thank Corette Wierenga for providing organotypic hippocampal slices. We want to thank Anna Akhmanova, Lukas Kapitein, and Corette Wierenga for critical reading of the manuscript. This work was supported by the Netherlands Organization for Scientific Research (Graduate Program of Quantitative Biology and Computational Life Sciences to NS), the European Research Council (ERC-StG 716011), and the Brain and Behavior Research Foundation (NARSAD Young Investigator Award) to HDM.

## AUTHOR CONTRIBUTIONS

Conceptualization, Methodology, Validation and Formal Analysis, J.W., A.P.H.J, N.S. and H.D.M., Investigation, J.W., A.P.H.J, N.S. and H.D.M., Resources, H.D.M. Writing - Original Draft & Editing, J.W., A.P.H.J, N.S. and H.D.M., Visualization, J.W., A.P.H.J, N.S. and H.D.M., Supervision, H.D.M., Funding Acquisition, H.D.M.

## DECLARATION OF INTRESTS

The authors declare no competing interest

## MATERIALS AND METHODS

### Antibodies and reagents

Primary antibodies used in this study are the following: rabbit anti-GFP (MBL Sanbio, 598, RRID AB_591819), mouse anti-PSD95 antibody ([K28/43], Neuromab, 75-028, RRID AB_2307331), mouse anti-alpha-tubulin ([B-5-1-2], Sigma, T5168, RRID AB_477582) and ATTO647N-conjugated anti-GFP nanobodies (GFPBooster-ATTO647N, Chromotek). Alexa488, Alexa568, Alexa594 and Alexa647 conjugated secondary antibodies were from Life Technologies. ATTO647n conjugated secondary antibodies were from Sigma. Phalloidin-conjugated A594 was from Life Technologies.

### Ethics statement

All experiments were approved by the Dutch Animal Experiments Committee (Dier Experimenten Commissie [DEC]), performed in line with institutional guidelines of Utrecht University, and conducted in agreement with Dutch law (Wet op de Dierproeven, 1996) and European regulations (Directive 2010/63/EU). Timed pregnant Wistar rats were obtained from Janvier. Wildtype male and female mice were used.

### Dissociated neuronal cultures

Dissociated hippocampal and cortical cultures were prepared from embryonic day 18 (E18) rat brains of both genders, as described in (60). Dissociated neurons were plated on Ø18mm coverslips coated with poly-L-lysine (37.5 µg/ml, Sigma-Aldrich) and laminin (1.25 µg/ml, Roche Diagnostics) at a density of 100,000 neurons per well. Cultures were grown in Neurobasal medium (NB), supplemented with 1% penicillin and streptomycin (pen/strep), 2% B27, and L-glutamine (all from Gibco) (NB-complete medium) at 37**°**C in 5% CO_2_. From DIV 1 onwards, medium was refreshed weekly by replacing half of the medium with Brainphys neuronal medium supplemented with 2% NeuroCult SM1 neuronal supplement (Stem cell technologies) and 1% pen/strep (BP-complete medium).

### Organotypic hippocampal slice cultures

Organotypic hippocampal slice cultures were prepared from wildtype mice at postnatal day 6-8 (adapted from Stoppini et al., 1991). After decapitation, the brain was quickly removed and placed in ice cold Gey’s Balanced Salt Solution (GBSS, containing (mM): 137 NaCl. 5 KCl, 1.5 CaCl2, 1 MgCl2, 0.3 MgSO4, 0.2 KH2PO4, 0.85 Na2HPO4) supplemented with 12.5 mM HEPES, 25 mM glucose and 1 mM kynurenic acid (pH set at 7.2, osmolarity set at 320 mOsm, sterile filtered). The frontal part of the brain and the cerebellum were removed along the transverse plane, and the hemispheres were then separated along the midline. Hippocampi were dissected and sliced perpendicularly to the long axis of the hippocampus with a thickness of 400 µm using a McIlwain Tissue Chopper. Slices were washed in culturing medium (consisting of 48% MEM, 25% HBSS, 25% horse serum, 30 mM glucose and 12.5 mM HEPES, pH set at 7.3-7.4 and osmolarity set at 325 mOsm) before being placed on Millicell cell culture inserts (Millipore) in 6-well plates containing culturing medium. Slices were kept at 37 °C with 5% CO_2_ until use and culturing medium was completely replaced twice a week.

## METHOD DETAILS

### Design and generation of ORANGE knock-in plasmids

#### Cloning of CRISPR/Cas9 knock-in vector pORANGE

To facilitate the generation of knock-in constructs, we developed a simple template vector (pORANGE). For this, we used pSpCas9(BB)-2A-Puro (PX459) V2.0 (Addgene 62988) and replaced SpCas9puro by SpCas9 from pAAV-nEFCas9 (Addgene 87115) flanked by the bipartite SV40 nuclear localization signal (NLS) sequences using the AgeI and EcoRI restriction sites, generating pSpCas9. To facilitate cloning of donor sequences, a multiple cloning site was inserted by annealing two complementary DNA oligos and ligation into the XbaI site of pSpCas9 generating pORANGE.

#### Design and cloning of ORANGE knock-in constructs

To select regions within a protein of interest suitable for introducing a tag, we carefully examined known protein functions, domains, presence of signal peptides, binding ligands and if known, protein structure, to minimize potential effects of the inserted tag sequence on protein function. For most proteins, this resulted in tagging close to the start or stop codon, or just behind the signal peptide. PAM sites in these identified regions were located in genomic sequences downloaded from the RGSC5.0/rn5 genome assembly through the UCSC genome browser gateway (https://genome-euro.ucsc.edu/). Target sequences were then chosen taking into consideration the MIT-guide specificity score (61) and Doench on-target scores (62). For some of the knock-ins, an extra G nucleotide was incorporated at the start of the target sequence to enhance transcription from the U6 promotor. We have no indication that this altered knock-in efficiency (For all protein target sites, target sequences and gRNA scores: see Table S1).

Next, oligos containing the 20-bp target sequences were annealed and ligated into the BbsI sites of pORANGE (Figure 1, S1B,C). Donor sequences were designed to contain the fluorescent tag sequence (GFP or mEos3.2) flanked by two Cas9 target sites identical to the genomic target site. Importantly, to facilitate genomic integration of the donor sequence in the correct orientation, these target sites including PAM sequences were inserted as the reverse complement of the genomic target sequence (Figure 1A, S1A). Additional linker sequences of at least three amino acids and additional base pairs to make the donor in frame after integration in the genome were introduced between the target sites and the tag sequence. Also, a start or stop codon was introduced in the donor sequence when proteins were tagged before the genomic start or stop codon respectively. Primer oligos with overhangs containing all these features where designed to generate the complete donor sequence by PCR. (See Figure S1B,C for two example designs). The donor sequences were PCR amplified using a GFP or mEos3.2-containing plasmid as template, and ligated into the multiple cloning site of the pORANGE vector containing the inserted target sequence to generate the complete knock-in construct. (For all primers used to generate the knock-in donor inserts, see Table S2).

For lentiviral applications, the ORANGE system was split in two plasmids. To generate pFUGW-Cas9, SpCas9 (from pAAV-nEFCas9) was ligated into the AgeI and EcoRI sites of FUGW (Addgene 14883). To generate the gRNA and donor containing lentiviral plasmid, first, mCherry-KASH amplified from pAAV-mTubb3 (Addgene 87116) was ligated into the BshTI and EcoRI sites of pFUGW-Cas9 replacing Cas9, yielding pFUGW-mCherry-KASH. Then, the U6 promotor, gRNA, and the donor sequence were amplified by PCR from the pORANGE construct, and inserted into the PacI site of pFUGW-mCherry-KASH using Gibson assembly (NEBuilder HiFi DNA assembly cloning kit).

### Transfection of dissociated hippocampal cultures

Neurons were transfected at DIV 3 (for knock-in) or DIV 14-18 (for overexpression) using Lipofectamine 2000 reagent (Invitrogen). Briefly, for one Ø18mm coverslip covered with 100,000 neurons, 1 - 2 µg DNA was mixed with 3.3 µl lipofectamine in 200 µl NB medium and incubated for 30 minutes at room temperature. Next, 500 µl conditioned medium was transferred to a new culture plate and replaced by 300 µl NB supplemented with 0.5 mM L-glutamine. The DNA mix was added to the neurons and incubated at 37°C and 5% CO_2_. After 90 - 120 minutes, neurons were transferred to the new culture plate with conditioned medium and 500 µl new NB medium supplemented with L-glutamine, B27 and pen/strep, and kept at 37°C and 5% CO_2_ for at least three days (for overexpression) and between 1 – 18 days for knock-in depending on the experiment.

### Electroporation of dissociated cortical cultures

For electroporation, cortical neurons were collected directly after dissection and dissociation, in a 15 ml tube and centrifuged for 5 minutes at 200 × g. Neurons were re-suspended in AMAXA transfection solution (Lonza) (1.3 × 10^6^ neurons per sample), mixed with 4 µg DNA, transferred to a gene pulser cuvette (Biorad), and electroporated using a Lonza Nucleofector 2b. Immediately after electroporation, fresh 37°C NB medium supplemented with B27, L-glutamine and pen/strep, was added to the cuvette after which the neurons were plated on a coated Ø18mm coverslip using a Pasteur pipette. Neurons were incubated at 37°C and 5% CO_2_ for 3 hours after which all medium was replaced with fresh NB medium supplemented with B27, L-glutamine and pen/strep.

### Lentivirus production and infection

For lentivirus production, HEK293T cells were maintained at high growth rate in DMEM supplemented with 10% bovine serum and 1% pen/strep. One day after plating, cells were transfected using polyethylenimine (PEI, Polysciences) with 2^nd^ generation lentiviral packaging plasmids (psPAX2 and 2MD2.G) and a pFUGW construct containing the desired insert, at 1:1:1 molar ratio. 6 hours after transfection, cells were washed once with phosphate buffered saline (PBS) (Lonza), and medium was replaced with DMEM containing 1% pen/strep. 48 hours after transfection, the supernatant was harvested and briefly centrifuged at 700 × g to remove cell debris. The supernatant was concentrated using Amicon Ultra 15 100K MWCO columns (Milipore), Cas9 and knock-in viruses were mixed at 1:1 and used immediately for infection. For cultured hippocampal neurons at DIV 2-4, 2 to 4 µL virus was added per well, and neurons were fixed at DIV 21-23 with 4% PFA with sucrose for 10 minutes. For organotypic hippocampal slices, virus was injected into the CA1 region at DIV1 using an Eppendorf Femtojet injector. Slices were fixed at DIV10 with 4% PFA (w/v) paraformaldehyde in PBS for 30 minutes, washed 3 times 10 minutes with PBS and mounted with Vectashield (Vector Laboratories).

### Genomic DNA extraction and sequence analysis

Genomic DNA was isolated from electroporated neurons at DIV4. Neurons were lysed in lysis buffer (100 mM Tris, 50 mM EDTA, 40 mM NaCl, 0.2% SDS, pH 8.5) and incubated with 100 µg/ml Proteinase K (Roche) at 55°C for 2 hours, followed by 1 hour at 85°C to inactivate proteinase K. DNA was isolated by ethanol precipitation and dissolved in Qiagen elution buffer (10 mM Tris, pH 8.0). Genomic PCR was performed to amplify the 5’ and 3’ junctions of the integrated donor (for PCR primers used, see: Table S3) using a touchdown PCR and Phusion HF polymerase (Thermo Fisher Scientific). PCR products were separated using agarose gel electrophoresis and subsequently purified using a gel extraction kit (Qiagen). Purified PCR products were ligated into the pJET vector according to the manufacture protocol (pJET cloning kit, Thermo Fisher Scientific). Individual clones were analyzed by Sanger sequencing (Macrogen) with primers in pJET. Clones were analyzed from at least two independent experiments.

### Immunohistochemistry

Hippocampal neurons were fixed using 4% (w/v) paraformaldehyde (PFA) and 4% (w/v) sucrose in phosphate buffered saline (PBS) (PFA/Suc) for 10 minutes at room temperature and washed three times in PBS containing 0.1 M glycine (PBS/Gly). Neurons were blocked and permeabilized in blocking buffer (10% (v/v) normal goat serum (NGS) (Abcam) in PBS/Gly with 0.1% (v/v) Triton X100) for 1 hour at 37°C. Next, coverslips were incubated with primary antibodies diluted in incubation buffer (5% (v/v) NGS in PBS/Gly with 0.1% (v/v) Triton X100) overnight at 4°C. Coverslips were washed three times 5 minutes with PBS/Gly and incubated with Alexa-conjugated goat anti-rabbit secondary antibodies, diluted 1:400 in incubation buffer for 1 hour at room temperature. Coverslips were washed three times 5 minutes in PBS/Gly, dipped in miliQ water (MQ) and mounted in Mowiol mounting medium (Sigma).

### Confocal imaging and analysis

Confocal images were acquired with a Zeiss LSM 700 with 63x N.A. 1.40 oil objective. A z-stack containing 7-12 planes at 0.56 µm interval was acquired, with 0.1 µm pixel size and maximum intensity projections were made for analysis and display. All image analysis was performed using FIJI software (63). Quantifications were performed in Excel 2016.

#### Quantification of knock-in efficiency

Hippocampal neurons were transfected at DIV 3, with a 1:1 ratio mixture of β-tubulin-GFP knock-in construct and pSM155-mCherry. Coverslips were fixed 24, 48, 72, 96, 120 and 144 hours after transfection using 4% PFA/Suc for 10 minutes at room temperature, washed three times with PBS/Gly and mounted in Mowiol mounting medium. mCherry-positive neurons were manually counted and scored as being tubulin-GFP positive or negative. At least 1,000 mCherry-positive neurons from two independent cultures were scored for each time point.

#### Quantification of synaptic levels and enrichment of PSD95

Hippocampal neurons were transfected at DIV 3 with the PSD95-GFP knock-in construct (targeting PSD95), or at DIV 15 with PSD95-GFP overexpression plasmid or pSM155-GFP. At DIV 21, neurons were fixed and stained with mouse anti-PSD95 antibody 1:200 and Alexa594-conjugated secondary antibodies as described above. For each neuron, 50 circular regions of interest (ROI) with 1 µm diameter were drawn around PSD95-GFP positive synapses. For each ROI, the mean intensity of the GFP signal and anti-PSD95 staining was measured, background subtracted, and normalized to the mean intensity value of all ROIs for both individual channels. Normalized intensity values for the PSD95-GFP knock-in signal and anti-PSD95 signal of individual synapses were plotted. In total, 550 synapses from 11 neurons divided over two independent neuronal cultures were used in the quantification.

To determine relative synaptic PSD95 content, PSD95 staining intensity in 22 circular ROIs with 1 µm diameter around synapses per transfected (knock-in, overexpression or GFP control) neuron were measured. Similarly, an equal number of ROIs was drawn around PSD95 puncta of nearby, non-transfected neurons within the same image. Intensities of the anti-PSD95 channel were measured, and background subtracted. Relative PSD95 content was quantified as the average anti-PSD95 intensity in synapses of a transfected neuron divided by those of the non-transfected neurons. To measure synapse size, the GFP signal (for PSD95-GFP knock-in and overexpression neurons) or anti-PSD95 signal (for GFP control) was thresholded and individual synapses were detected using FIJI “Analyze Particles” with a detection size of 0.04-Infinity (µm^2^) with a detection circularity of 0-1. Measured values were plotted as averages per analyzed neuron. To analyze synaptic enrichment of PSD95, circular ROIs were drawn within synapses and on the dendritic shaft. Mean GFP intensity was measured, background subtracted and averaged per neuron. Plotted ratio is the average intensity of synaptic GFP signal divided by that of the dendritic shaft. For each condition, at least 15 neurons from two independent neuronal cultures were analyzed.

To compare PSD95 levels in transfected, but knock-in negative neurons, neurons were transfected with a 1:1 ratio of pHomer1c-mCherry and the pORANGE empty vector, or pHomer1c-mCherry and pPSD95-GFP knock-in construct at DIV 3. At DIV 21, neurons were fixed and stained for endogenous PSD95 as described above. Homer1c-mCherry positive neurons were used to locate transfected neurons, and to draw ROIs around synapses. For both conditions, 20 neurons from two independent neuronal cultures were analyzed.

#### Quantification of F-actin levels

Neurons were transfected with pGFP-β-actin knock-in construct at DIV 3. The pGFP-β-actin expression or pSM155-GFP overexpression plasmids were transfected at DIV 15. Neurons were fixed at DIV 21 with 4% PFA/Suc for 10 minutes at room temperature. Coverslips were washed three times 5 minutes with PBS/Gly, and blocked and permeabilized in blocking buffer (10% (v/v) NGS and 0.1% (v/v) Triton X100 in PBS/Gly) for 1 hour at 37°C. Next, the neurons stained with Phalloidin-Alexa594 (Invitrogen) diluted 1:200 in blocking buffer for 1 hour at room temperature. Coverslips were washed three times 5 minutes in PBS/Gly and mounted in Mowiol mounting medium. Per neuron, 50 circular ROIs of 1 µm in diameter manually drawn around spines. Similarly, an equal number of ROIs was drawn around puncta of nearby, non-transfected neurons within the same image. Mean intensities of GFP signal and phalloidin staining were measured. To correlate GFP-β-actin knock-in signal to the phalloidin staining, intensities of individual ROIs of transfected neurons were normalized to the average value for both the GFP signal and the phalloidin staining of all ROIs. Normalized values of individual spines were plotted. To measure relative F-actin levels in spines, phalloidin intensities of individual ROI measurements were background subtracted and averaged for each neuron. The average phalloidin intensity in the transfected neuron relative to the non-transfected neuron from the same image is plotted. In total, 9 neurons were used for analysis.

#### Quantification of NMDA receptor levels in synapses

Neurons were transfected at DIV 3 with a construct for GFP-GluN1 knock-in. Neurons were fixed at DIV 21 and stained with anti-PSD95 as described above. For each neuron, 50 circular ROIs with 1 µm diameter were drawn around GFP-GluN1 positive synapses. For each ROI, the mean intensity of the GFP signal and anti-PSD95 staining was measured, background subtracted, and normalized to the mean intensity value of all ROIs for both individual channels. Normalized intensity values for the GFP-GluN1 knock-in signal and anti-PSD95 signal of individual synapses were plotted. In total, 450 synapses from 9 neurons divided over two independent neuronal cultures were used in the quantification.

### gSTED imaging and analysis

Hippocampal neurons were transfected with indicated knock-in constructs at DIV 3 and fixed at DIV 21. Dual-color gated STED (gSTED) imaging was performed on PSD95-GFP, GFP-β-actin GFP-GluN1 and GFP-CaMKIIα knock-in neurons stained with anti-GFP and anti-PSD95. (Dual-color) gSTED imaging was additionally performed on extracted cytoskeleton of the GFP-β-actin and β3-tubulin-GFP knock-in neurons. At DIV 7 (β3-tubulin-GFP knock-in) and DIV 21 (GFP-β-actin knock-in), the neuronal cytoskeleton was extracted using extraction buffer (PEM80-buffer (80 mM PIPES, 1 mM EGTA, 2 mM MgCl2, pH 6.9), 0.3% Triton-X, 0.1% glutaraldehyde) for 1 minute at room temperature. Next, neurons were fixed with PFA/Suc for 10 min at room temperature, washed three times 5 minutes with PBS/Gly, and subsequently incubated with 1 mg/ml sodium borohydride in PBS for 7 minutes at room temperature. Coverslips were washed 3 times 5 minutes with PBS/Gly. The GFP signal was additionally stained with anti-GFP. The β3-tubulin-GFP knock-in was additionally stained for alpha-tubulin, diluted 1:1000. Anti-GFP primary antibodies were always stained with ATTO647N-conjugated secondary antibody, and anti-PSD95 and anti-alpha-tubulin were stained with the Alexa594-conjugated secondary antibody. To label surface receptors, GFP-GluN1 knockin neurons were stained with anti-GFP prior to permeabilization, and anti-PSD95 staining. Imaging was performed with a Leica TCS SP8 STED 3X microscope using a HC PL APO 100x/ N.A. 1.4 oil immersion STED WHITE objective. The 590 nm and 647 nm wavelengths of pulsed white laser (80MHz) were used to excite the Alexa594-labeled and the ATTO647N-labeled proteins, respectively. Both Alexa594 and ATTO647N were depleted with the 775 nm pulsed depletion laser (50-75% of maximum power) and we used an internal Leica HyD hybrid detector (set at 100% gain) with a time gate of 0.3 ≤ tg ≤ 6 ns. Multiple Z-stack were obtained at 0.2 µm interval to acquire image stacks in 2D STED mode using the 100x objective. Maximum intensity projections were obtained for image display and analysis.

#### Quantification of colocalization

Using ImageJ software a line scan of interest was drawn to obtain pixel intensity data to assess the degree of colocalization between two structures along that line. To quantify the degree of colocalization between two structures, entire images, showing parts of the dendritic tree of a knock-in neuron, were used for analysis. First, all dendritic spines (positive for both proteins; PSD95-GFP knock-in and anti-PSD95 staining or GFP-β-actin knock-in and anti-PSD95 staining) were selected by drawing ROIs in ImageJ. Next, the ROIs were combined to clear the outside of the ROIs to remove all background from surrounding neurons or dendritic shafts. Then, the ImageJ plugin “JaCoP” (Just Another Colocalization Plugin) was used to calculate the Pearson’s correlation coefficient (PCC) and Manders’ overlap coefficient (MOC). For the MOC the thresholding was done manually. These analyses were performed on both the confocal and STED maximum projections of the exact same regions (of a neuron). In total, 10 PSD95-GFP knock-in and 7 GFP-β-actin knock-in neurons were analyzed from two independent experiments.

#### Quantification of FWHM GluN1 substructures

The FIJI plugin FWHM (Full width at Half Maximum) macro developed by John Lim was used to measure the FWHM from intensity profiles using Gaussian fitting. Line scans were drawn along the width and length of identified GluN1 substructures (by setting an appropriate brightness/contrast) to obtain the FWHM of the length and width of these substructures. Subsequently, these substructures were categorized as synaptic or extra-synaptic, based on the colocalization with PSD95. For image display, the length was plotted against the width for each cluster. 479 GFP-GluN1 clusters (387 synaptic, 92 extra-synaptic) from 3 neurons were analyzed.

#### Quantification of total GluN1 cluster area and number of substructures per PSD

For the quantification of total GluN1 cluster area per PSD, and correlation with PSD area, the same images were used as for the quantification of the FWHM of the GluN1 substructures. Specifically, the STED resolved images were used for the quantification of GluN1 cluster area, whereas the confocal images were used to quantify the area of the PSD, using PSD95 as a marker. First, an ROI was drawn around the knock-in neuron of interest, to clear the outside of the ROI removing all background. Subsequently, the image was subjected to thresholding to isolate the objects of interest from the background and Watershedding to separate overlapping objects. Then, all objects (GluN1 clusters and PSDs) were detected using “Analyze Particles” with a detection size of 0.02-Infinity (µm^2^) for GluN1 substructures and 0.04-Infinity (µm^2^) for PSDs and all with a detection circularity of 0-1. For this analysis only synaptic GluN1 substructures were used. If multiple GluN1 substructures were detected per PSD, the areas were added up to a total GluN1 cluster area per PSD. For image display, the total GluN1 cluster area was plotted against PSD area, together with a nonlinear regression curve fit. Additionally, the number of GluN1 substructures detected per PSD were plotted.

### Single-molecule localization microscopy and detection

dSTORM imaging was performed on a Nikon Ti microscope equipped with a Nikon 100x N.A. 1.49 Apo TIRF oil objective, a Perfect Focus System. Effective pixel size is 65 nm. Oblique laser illumination was achieved using a custom illumination pathway with a 60 mW 405 nm diode laser (Omicron), a 200 mW 491 nm diode laser (Omicron) and a 140 mW 641 nm diode laser (Omicron). Emission light was separated from excitation light with a quad-band polychroic mirror (ZT405/488/561/640rpc, Chroma), and an additional band-pass emission filters (ET 525/595/700, Chroma). Fluorescence emission was acquired using an ORCA-Flash 4.0v2 CMOS camera (Hamamatsu). Lasers were controlled using Omicron software while all other components were controlled by μManager software (64).

Live-cell SMLM imaging experiments were performed on a Nikon Ti microscope equipped with a 100× N.A. 1.49 Apo TIRF oil objective, a Perfect Focus System and an additional 2.5× Optovar to achieve an effective pixel size of 64 nm. Oblique laser illumination was achieved using a custom illumination pathway with a AA Acousto-optic tunable filter (AA opto-electronics), 15 mW 405 nm diode laser (Power Technology), a 100 mW 561 nm DPSS laser (Cobolt Jive), and a 40 mW 640 nm diode laser (Power Technology). Emission light was separated from excitation light with a quad-band polychroic mirror (ZT405/488/561/640rpc, Chroma), and an additional band-pass emission filters (ET 525/595/700, Chroma). Fluorescence emission was acquired using a DU-897D EMCCD camera (Andor). All components were controlled by μManager software (64).

Acquired image stacks were analyzed using the ImageJ plugin DoM (Detection of Molecules) v1.1.5 (65). Briefly, each image was convoluted with a 2D Mexican hat-type kernel that matches the microscope’s point spread function. Spots were detected by thresholding the images and localized by fitting a 2D Gaussian function using unweighted nonlinear least-squares fitting with the Levenberg-Marquardt algorithm. Drift correction was applied by calculating the spatial cross-correlation function between intermediate super-resolved reconstructions.

#### Single-molecule tracking PALM and analysis

Hippocampal neurons were transfected with pmEos3.2-CaMKIIα knock-in construct at DIV 3 and imaged at DIV 21–23. Neurons were imaged in extracellular imaging buffer (10 mM HEPES, 120 mM NaCl, 3 mM KCl, 2 mM CaCl_2_, 2 mM MgCl_2_, 10 mM glucose, pH 7.35) at RT. mEos3.2 molecules were photoconverted from green to red fluorescence using simultaneous 405 nm and 561 nm illumination using total internal reflection fluorescence (TIRF). Stacks of 5,000 – 7,000 frames were acquired at 50 Hz. PALM reconstruction was made in DoM, plotting localizations based on their localization precision, rendered with a pixel size of 10 × 10 nm. Molecules localized with precision < 25 nm were used for further analysis. Tracking was accomplished using custom tracking algorithms in MATLAB (MathWorks) using a tracking radius of 512 nm. For tracks consisting of ≥ 4 frames the instantaneous diffusion coefficient was estimated as described (38). The first three points of the mean squared displacement (MSD) vs. elapsed time (*t*) plot were used to fit the slope using linear fitting adding a value of 0 at MSD(0). Tracks with a negative slope (< 8%) were ignored. The diffusion coefficient *Deff* was then calculated using MSD = *4Deff t*. Individual tracks were plotted using MATLAB, each given a random color. All single-molecule trajectories from all acquisitions were used to visualize a frequency distribution. On this, we fitted two gaussian distributions to identify the two kinetic populations. Mean values for the two fits were calculated per analyzed cell and plotted. In total, 11 neurons, from two independent experiments we included in the analysis.

#### dSTORM imaging and analysis

Hippocampal neurons were transfected at DIV 3 with the pGFP-GluN1 knock-in construct and fixed on DIV 21. Neurons were surface stained with anti-GFP 1:2000 and Alexa647-conjugated secondaries as described above. Neurons were post-fixed in 4% PFA/Suc for 5 minutes, additionally washed 3 times with PBS/Gly. and kept in PBS at 4°C until imaging. dSTORM imaging was performed in PBS containing 10 - 50 mM MEA, 5% w/v glucose, 700 μg/ml glucose oxidase, and 40 μg/ml catalase. GFP-GluN1 knock-in positive neurons were located on GFP signal. For dSTORM, the sample was illuminated (in TIRF) with continuous 647 nm laser light and gradually increasing intensity of 405 nm laser light. Stacks of 10,000 – 15,000 frames were acquired at 50 Hz. dSTORM reconstruction was made in DoM, plotting localizations based on their localization precision, rendered with a pixel size of 10 × 10 nm. Molecules with a localization precision < 15 nm were selected for further analysis. Next, blinking events longer than one frame, were filtered out by tracking (tracking radius of 130 nm). GluN1 clusters were identified using the DBSCAN algorithm (48) implemented in MATLAB. Subsequently, the alpha shape was used as the cluster border. Clusters with a density of > 5000 molecules/µm were used for further analysis. For each individual cluster, molecules were plotted and color-coded according to the local density (42), defined as the number of molecules within a radius of 5 times the mean nearest neighbor distance of all molecules within the cluster. Molecules with a local density value > 40 were considered to be enriched in a nanodomain. Nanodomains were isolated using MATLAB functions linkage() and cluster(). The polygon circumventing molecules belonging to individual nanodomains was used to calculate the diameter of the nanodomain. Nanodomains containing < 5 localizations and diameter < 30 nm were rejected. In total, 859 clusters from 3 neurons, from two independent experiments were analyzed.

#### uPAINT and analysis

Neurons transfected with the pGFP-GluN1 knock-in construct were imaged at DIV 21-23 in extracellular imaging buffer supplemented with 0.8% BSA. GFP-GluN1 positive neurons were identified by GFP signal and ATTO647N-conjugated anti-GFP nanobodies (GFPBooster-ATTO647N, Chromotek) were bath applied to a final dilution of 1:50,000. Imaging was conducted at 50 Hz frame rate with 640 nm excitation laser illumination (in TIRF). Molecules fitted with a precision < 50 were tracked with tracking radius of 512 nm and diffusion coefficient determined for tracks > 30 frames. A cell mask was drawn manually to filter out localizations outside neurons due to non-specifically bound nanobody. Tracking and estimation of the instantaneous diffusion was performed as described for the PALM imaging. Synapses were identified based on widefield Homer1c-mCherry signal as described (66). Synaptic tracks were defined as tracks of which 80% of the localizations were located within the border of the synapse. All others were considered extrasynaptic. In total, 6 neurons from three independent experiments were analyzed.

### Statistics

Statistical significance was tested with a student’s t-test when comparing two groups (Figures 3E, 3G) A *p*-value below 0.05 was considered significant. If multiple groups were compared (Figures 3C, 3D, 5K, 5L, 5M, S4C), statistical significance was tested with a one-way ANOVA followed by a Bonferroni’s multiple comparison. In all figures * was used to indicate a *p*-value < 0.05, ** for *p* < 0.01, and *** for *p* < 0.001. Reported *n* is number of neurons, and each experiment was replicated in cultures from at least 2 independent preparations. Statistical analysis and graphs were prepared in GraphPad Prism and figures were generated in Adobe Illustrator CC.

## DATA AND SOFTWARE AVAILABILITY

Custom MATLAB routines are available upon request.

## ADDITIONAL RESOURCES

Plasmids from this study will be made available through Addgene.

---

**Figure S1 related to Figure 1.**
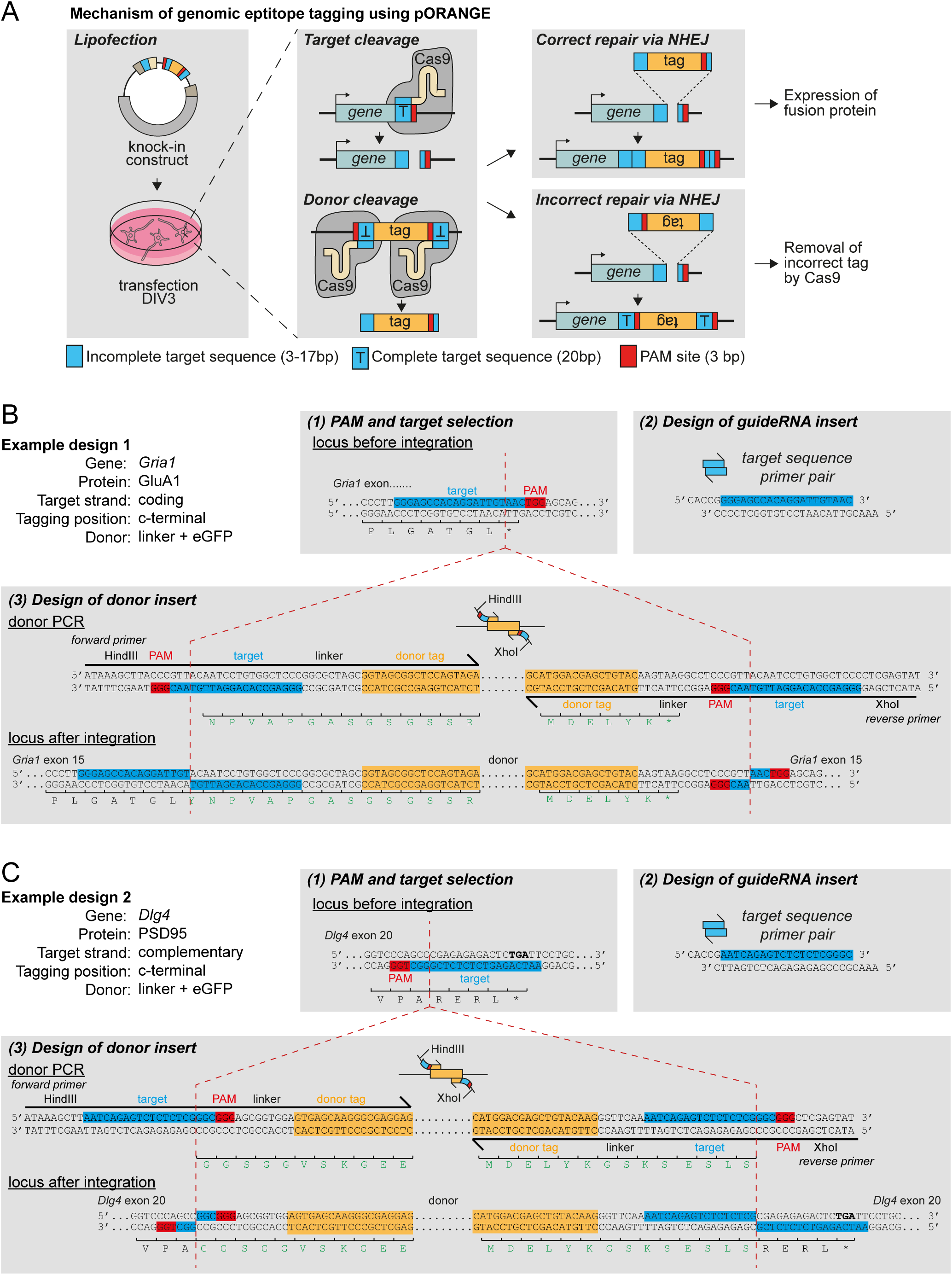
Mechanism of ORANGE knock-in and design of knock-in constructs. **(A)** Mechanism of ORANGE knock-in approach. **(B, C)** Examples of knock-in construct design for two genes with target sequences in opposite genomic strands. The target sequence is indicated in blue, the PAM sequence in red and the part of the primer used for PCR amplification of the donor DNA in shown in yellow. Amino-acid sequence is shown under the sequences. *, indicates stop codon. Red dotted lines indicate position of Cas9 cleavage and sites of integration.

**Figure S2 related to Figure 3.**
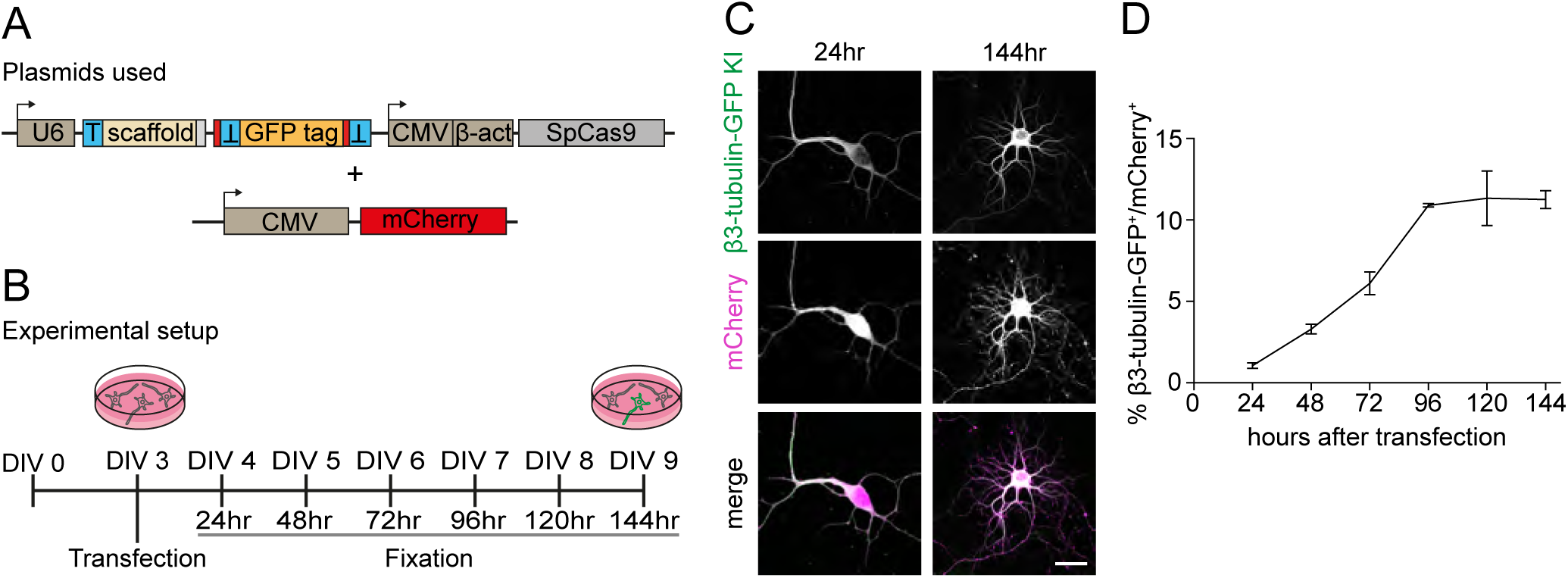
Efficiency of ORANGE knock-in over-time. **(A)** Schematic overview of knock-in and mCherry reporter plasmids, and **(B)** experimental setup. **(C)** Representative images of β3-tubulin-GFP knock-in (green) together with soluble mCherry fill (magenta), fixed 24 hours (DIV 4) and 144 hours (DIV 9) after transfection. Scale bar, 20 µm. **(D)** Quantification of β3-tubulin-GFP knock-in efficiency over time as percentage of transfected (mCherry positive) neurons. Data are represented as mean ± SEM.

**Figure S3 related to Figure 3.**
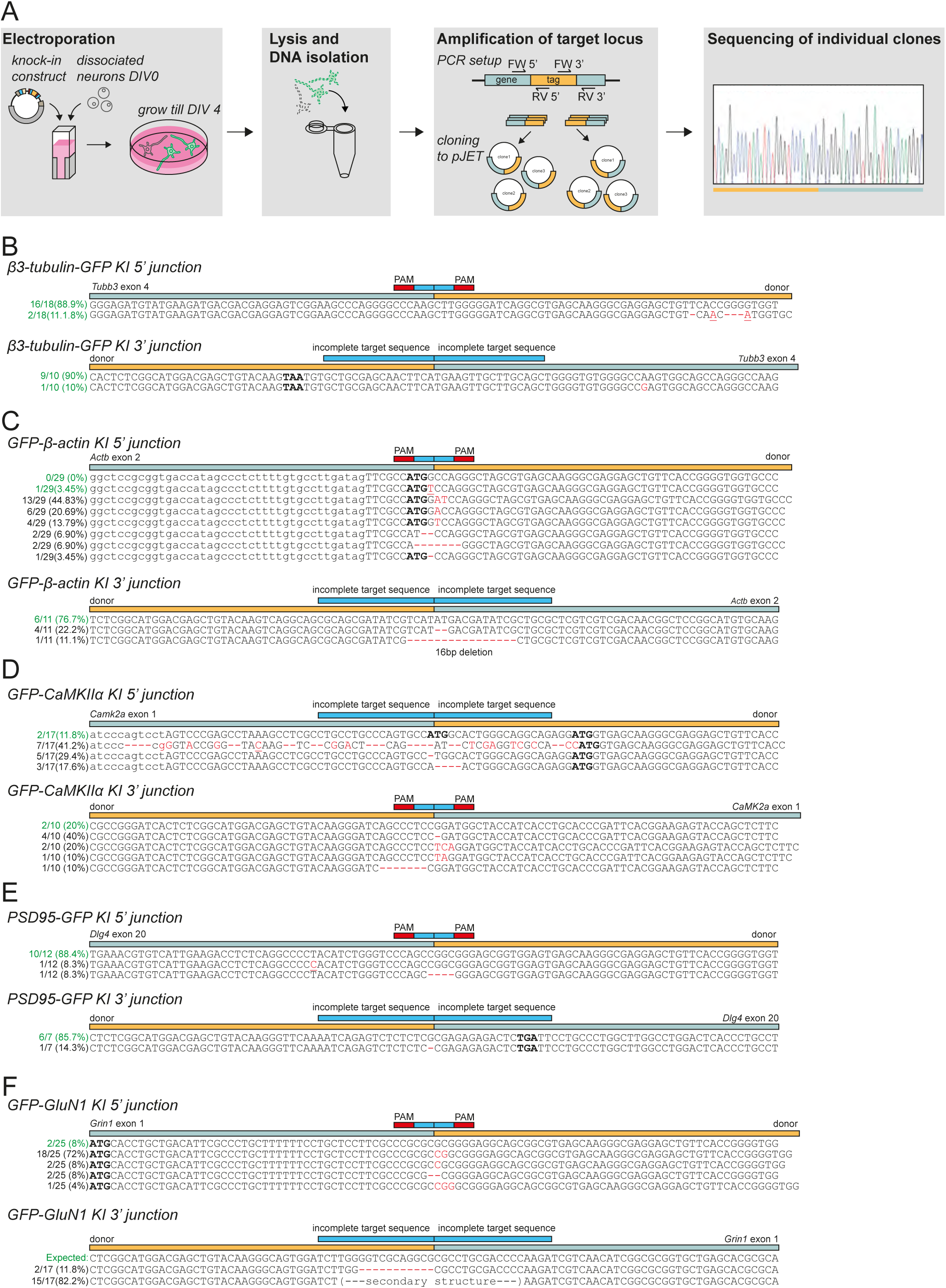
Genomic analysis of donor integration at targeted locus. **(A)** Experimental setup. Neurons were electroporated immediately after dissociation and grown until DIV 4. DNA was isolated, and the target locus was amplified, and cloned into pJET. Individual clones were then sequenced. **(B-F)** Sequencing analysis of individual pJET clones for selected knock-ins. Bars above sequences indicate genomic sequence (light-blue) and donor sequence (yellow). Number of clones and the percentage showing in-frame integration are shown in green. In the sequence, start and stop codons (if present) are highlighted in bold. Deletions are shown as red dashes. Indels are indicated in red. Point mutations are in red and underlined.

**Figure S4 related to Figure 5.**
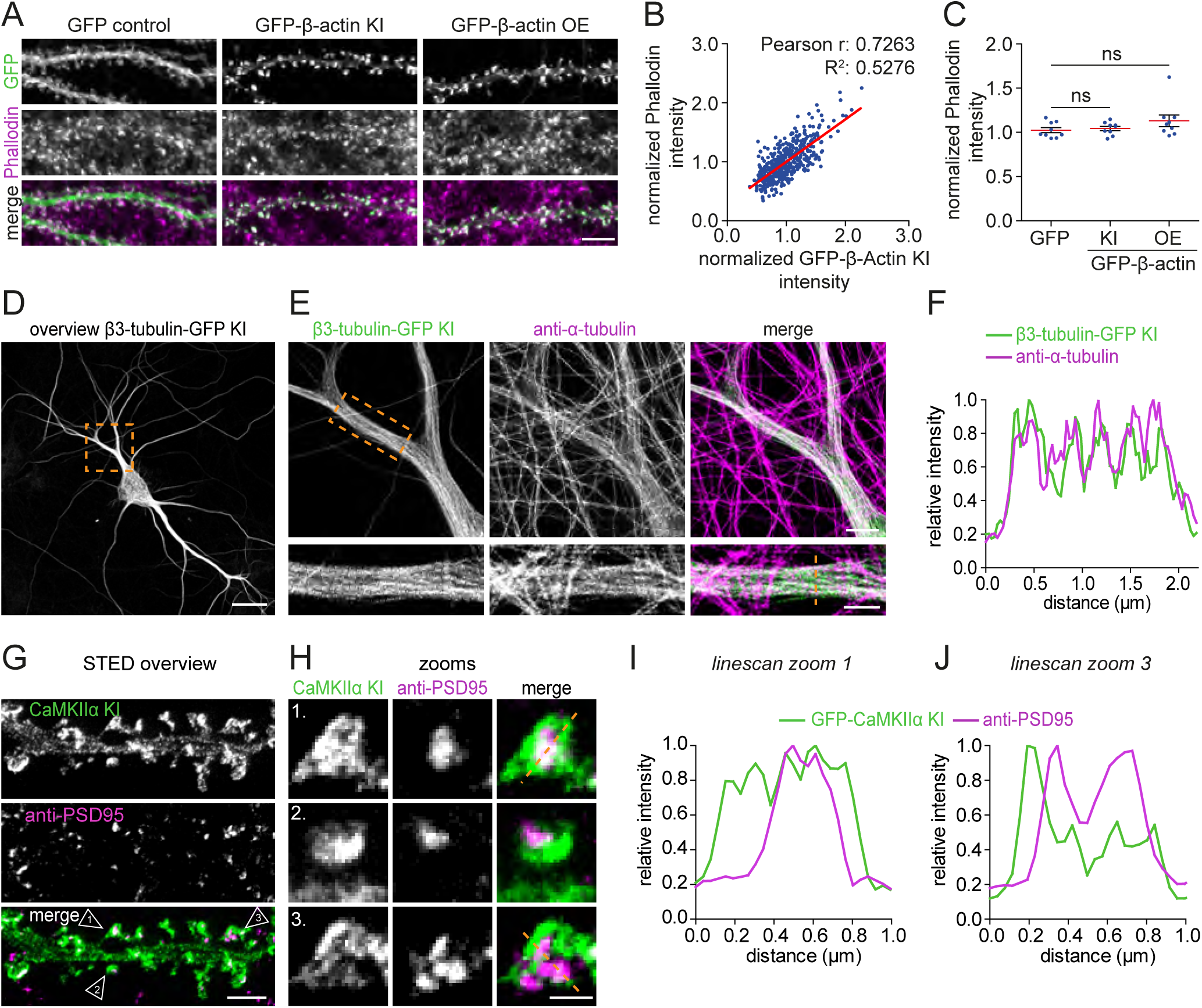
β-actin knock-in validation and dual-color gSTED imaging of endogenous β3-tubulin and CaMKIIα. **(A)** Representative images of dendrites expressing soluble GFP, GFP-β-actin knock-in or GFP-β-actin overexpression construct (green), stained with phalloidin (magenta). DIV 21. Scale bar, 5 µm. **(B)** Correlation between GFP-β-actin knock-in and phalloidin staining intensity within dendritic spines. **(C)** Quantification of F-actin levels compared to non-transfected neurons. **(D)** Representative gSTED overview image and zooms **(E)** of β3-tubulin-GFP knock-in extracted and stained with anti-GFP (ATTO647N; green), and anti-alpha-tubulin (Alexa594; magenta). DIV 7. Cytosolic monomeric tubulin was removed using an extraction protocol. Scale bars, 20 µm, 4 µm and 2 µm for the overview and zooms respectively. **(F)** Intensity profile along the line indicated in (E). **(G)** gSTED of dendrites and zooms **(H)** positive for GFP-CaMKIIα knock-in stained with anti-GFP (Atto647N; green) and anti-PSD95 (Alexa594; magenta). Scale bars, 2 µm and 500 nm for (G) and (H) respectively. **(I, J)** Line scans of individual spines indicated in (H). Data is represented as mean ± SEM. ns, indicates not significant. ANOVA.

**Figure S5 related to Figure 5.**
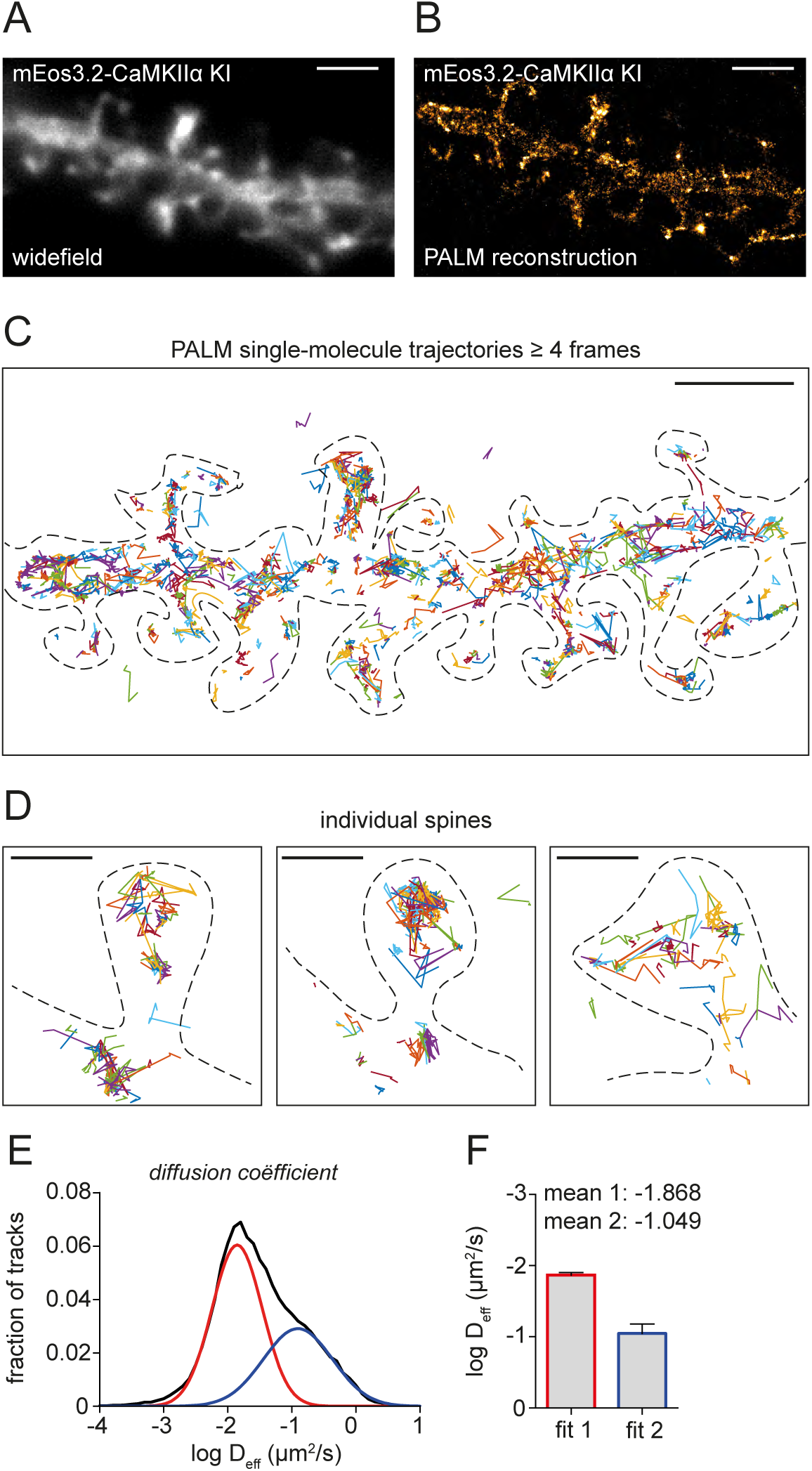
Live-cell super-resolution PALM imaging of endogenous CaMKIIα dynamics. **(A)** Example of dendrite expressing mEos3.2-CaMKIIα knock-in. Scale bar, 2 µm. **(B)** Single-molecule PALM reconstruction of dendrite shown in (A). DIV 21. Scale bar, 2 µm. **(C)** Individual single-molecule trajectories. Scale bar, 2 µm. Dotted line indicates cell outline. **(D)** Representative zooms of single-molecule trajectories in individual spines. Scale bar, 200 nm. **(E)** Frequency distribution of diffusion coefficients derived from single-molecule trajectories (black line). Two gaussians fits (red and blue) indicate two kinetic populations. **(F)** Quantification of mean diffusion coefficient for each of the two kinetic populations. Data is represented as mean ± SEM.

**Table S1.**
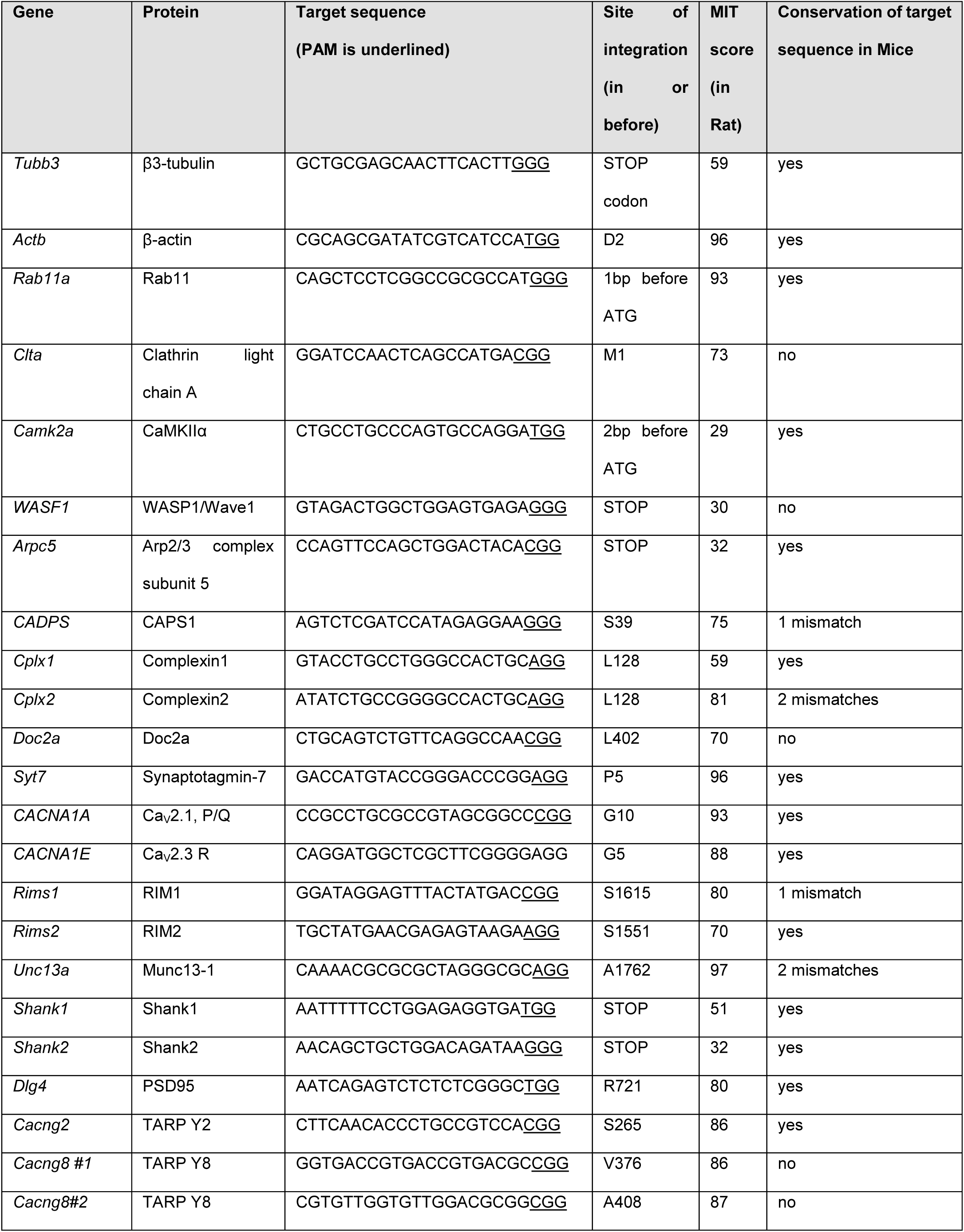
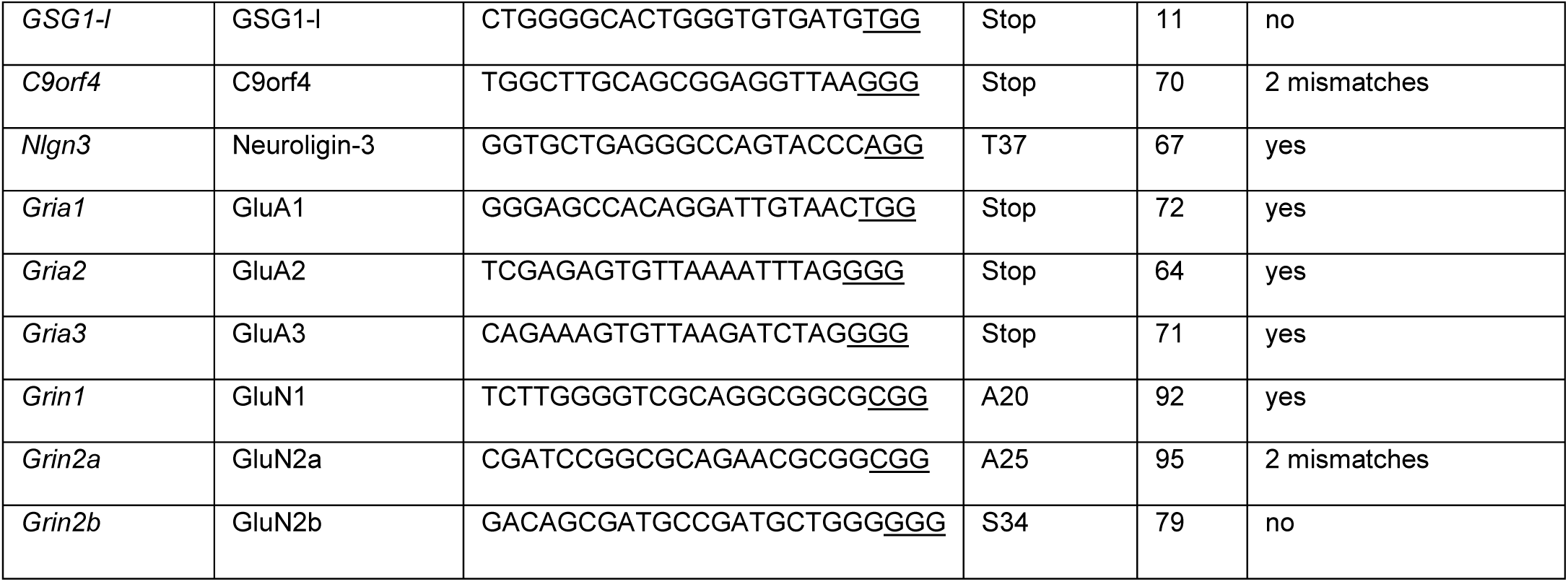
Overview of selected genes and target sequences for CRISPR/Cas9 knock-ins.

**Table S2.**
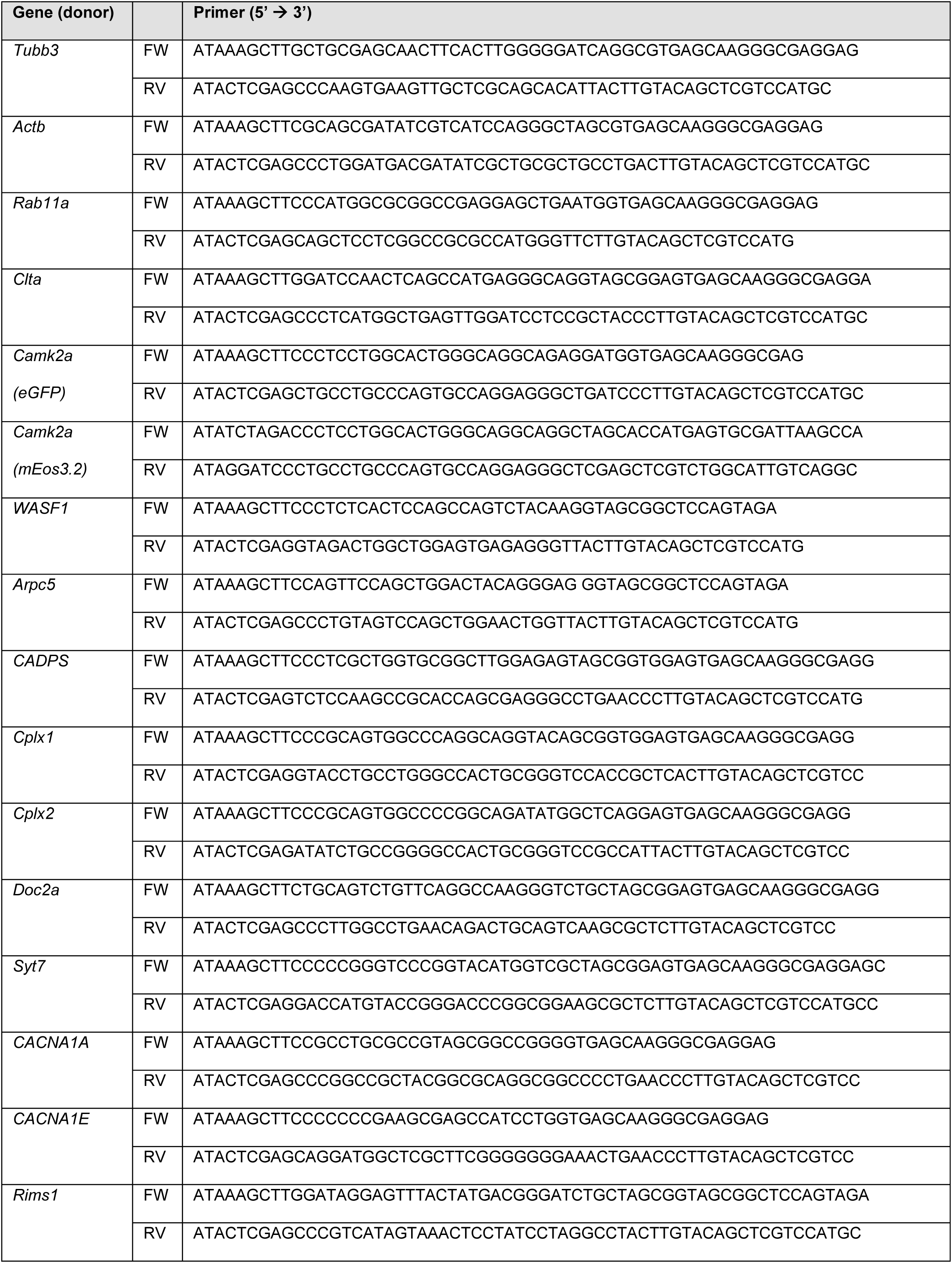
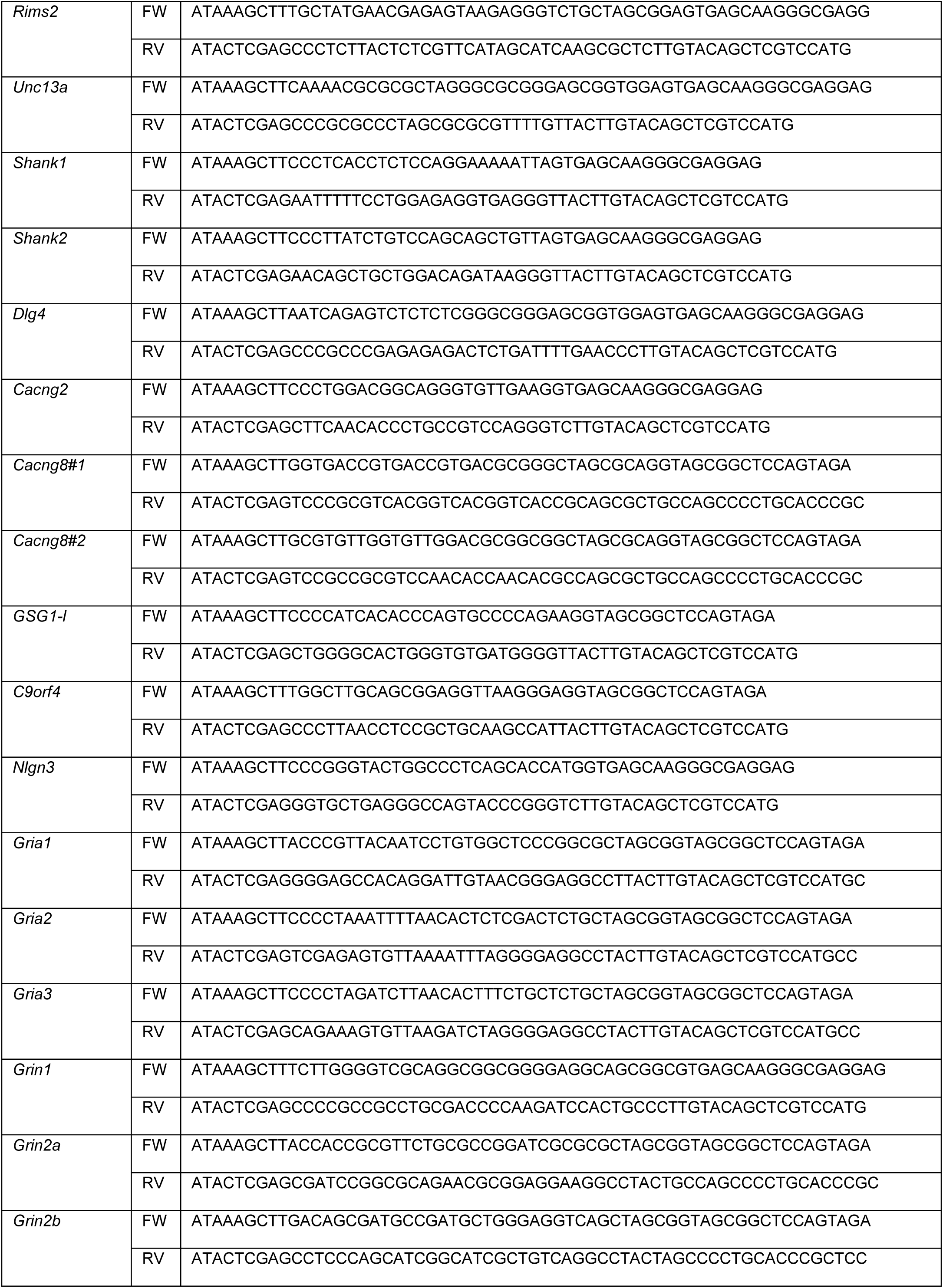
Donor PCR primers.

**Table S3.**
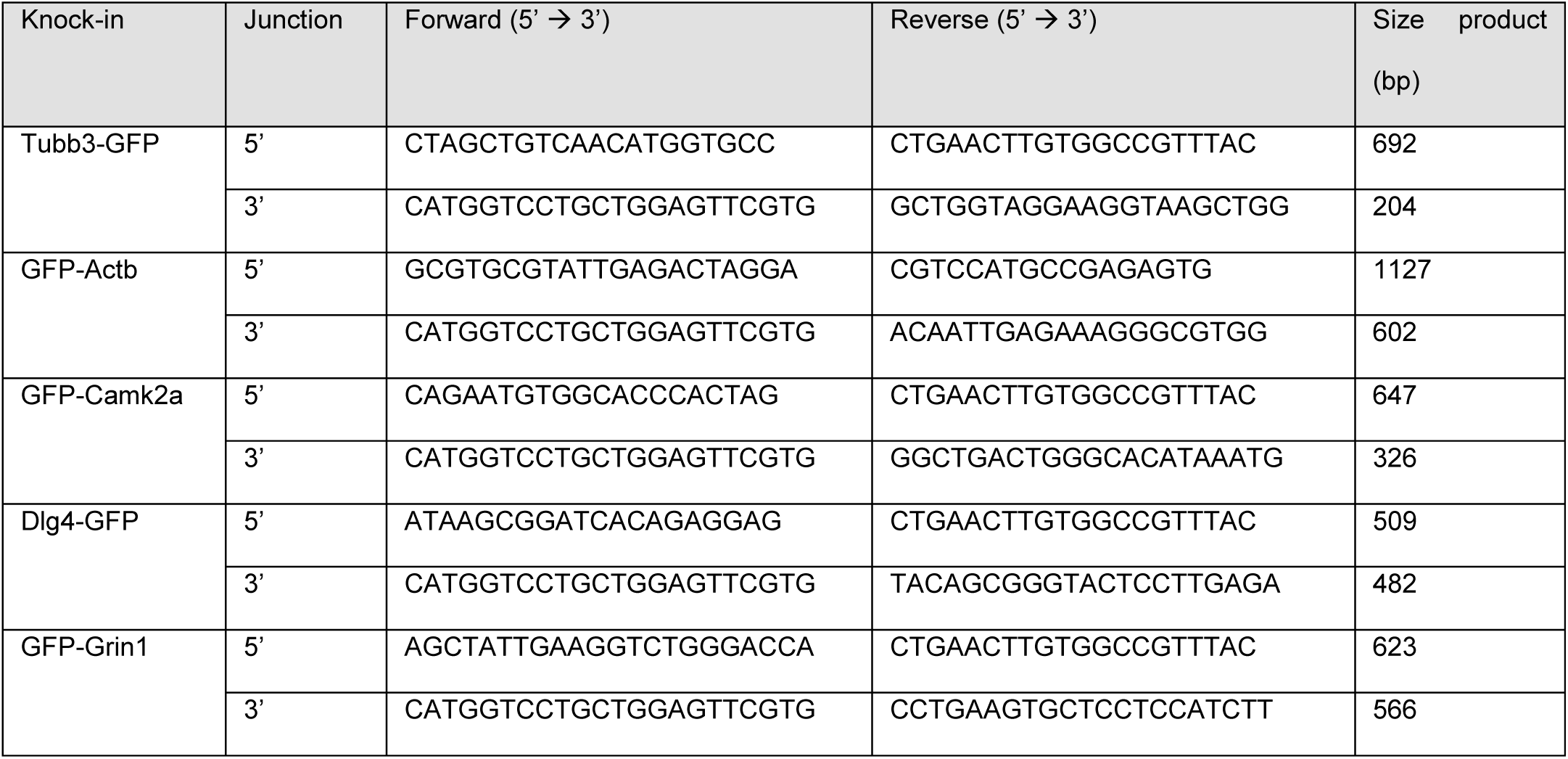
Genomic PCR primers.

